# Pharmacological inhibition of tyrosine protein-kinase 2 reduces islet inflammation and delays type 1 diabetes onset in mice

**DOI:** 10.1101/2024.03.20.585925

**Authors:** Farooq Syed, Olivia Ballew, Chih-Chun Lee, Jyoti Rana, Preethi Krishnan, Angela Castela, Staci A. Weaver, Namratha Shivani Chalasani, Sofia F. Thomaidou, Stephane Demine, Garrick Chang, Alexandra Coomans de Brachène, Maria Ines Alvelos, Lorella Marselli, Kara Orr, Jamie L. Felton, Jing Liu, Piero Marchetti, Arnaud Zaldumbide, Donalyn Scheuner, Decio L. Eizirik, Carmella Evans-Molina

## Abstract

Tyrosine protein-kinase 2 (TYK2), a member of the Janus kinase family, mediates inflammatory signaling through multiple cytokines, including interferon-α (IFNα), interleukin (IL)-12, and IL-23. Missense mutations in TYK2 are associated with protection against type 1 diabetes (T1D), and inhibition of TYK2 shows promise in the management of other autoimmune conditions. Here, we evaluated the effects of specific TYK2 inhibitors (TYK2is) in pre-clinical models of T1D. First, human β cells, cadaveric donor islets, and iPSC-derived islets were treated *in vitro* with IFNα in combination with a small molecule TYK2i (BMS-986165 or a related molecule BMS-986202). TYK2 inhibition prevented IFNα-induced β cell HLA class I up-regulation, endoplasmic reticulum stress, and chemokine production. In co-culture studies, pre-treatment of β cells with a TYK2i prevented IFNα-induced activation of T cells targeting an epitope of insulin. *In vivo* administration of BMS-986202 in two mouse models of T1D (*RIP-LCMV-GP* mice and NOD mice) reduced systemic and tissue-localized inflammation, prevented β cell death, and delayed T1D onset. Transcriptional phenotyping of pancreatic islets, pancreatic lymph nodes (PLN), and spleen during early disease pathogenesis highlighted a role for TYK2 inhibition in modulating signaling pathways associated with inflammation, translational control, stress signaling, secretory function, immunity, and diabetes. Additionally, TYK2i treatment changed the composition of innate and adaptive immune cell populations in the blood and disease target tissues, resulting in an immune phenotype with a diminished capacity for β cell destruction. Overall, these findings indicate that TYK2i has beneficial effects in both the immune and endocrine compartments in models of T1D, thus supporting a path forward for testing TYK2 inhibitors in human T1D.

## INTRODUCTION

During the development of type 1 diabetes (T1D), cells of the innate and adaptive immune system infiltrate the pancreatic islets, produce a variety of pro-inflammatory cytokines, and create a feed-forward cycle of immune activation, inflammation, and β cell death. Tyrosine protein-kinase 2 (TYK2), a member of the Janus kinase (JAK) family, transduces signals from type I and type II cytokine receptors via phosphorylation and activation of the transcription factors, signal transducer and activator of transcription 1 and 2 (STAT1 and 2) (1,2). Polymorphisms in TYK2 are associated with a reduced risk of several autoimmune diseases, including T1D, ulcerative colitis, and rheumatoid arthritis (3–5). Notably, a polymorphism causing a missense mutation in TYK2, leading to decreased function, is associated with protection against T1D (3,6,7).

TYK2 modulates signaling through three specific cytokines: interferon-α (IFNα), interleukin (IL)-12, and IL-23 (8,9). While each of these cytokines has been linked with T1D pathogenesis, clinical and preclinical data suggest a prominent role for IFNα in mediating inflammatory crosstalk between β cells and the immune system during T1D progression. There is evidence of a type 1 interferon transcriptional signature in the peripheral blood of children at high genetic risk for T1D even prior to seroconversion (10,11) and in the whole blood and pancreas of individuals with recent onset T1D (12). IFNα activates pancreatic islet-resident macrophages and T cells, while also impacting stress pathways in the β cell (13–16). Findings by our group and others have shown that treatment of human β cells with IFNα augments HLA class I overexpression, modulates transcriptional programs associated with endoplasmic reticulum (ER) stress and inflammatory signaling, and increases alternative RNA splicing and the diversity of transcripts expressed by β cells (17–21). Furthermore, case reports have documented T1D onset in conjunction with the therapeutic use of IFNα in hepatitis C and leukemia (22,23). In contrast, antibody-mediated blockade of IFNα or the type I IFN α/β receptor (IFNAR1) is sufficient to prevent the development of T1D in rodent models (24–27).

Blocking IFNα-mediated effects via inhibition of JAK tyrosine kinases has shown beneficial effects in *in vivo* murine and *in vitro* cell models of T1D (28,29). Additionally, in a recent clinical trial, the JAK1/2 inhibitor baricitinib preserved C-peptide in individuals 10-30 years of age who were newly diagnosed with Stage 3 T1D (30). Pharmacological inhibition of TYK2 may offer safety advantages over JAK inhibitors, which are associated with an increased risk of venous and arterial thrombotic events, cardiovascular events, and malignancy in older populations (31,32). In 2022, a selective allosteric inhibitor targeting the TYK2 pseudokinase domain (deucravacitinib; BMS-986165) was approved by the U.S. Food and Drug Administration (FDA) for use in psoriasis (33). Previous *in vitro* studies suggest TYK2 inhibitors are able to block IFNα-mediated signaling in human islets and β cell lines (34,35). In addition, a recent study showed that NOD mice with total body TYK2 deletion were protected from the development of T1D (36).

To define how pharmacological TYK2 inhibition mediates *in vivo* crosstalk between the immune system and the β cells, we tested the effects of two TYK2is, BMS-986165 and BMS-986202, in three *in vitro* human model systems and then determined the impact of BMS-986202 in two mouse models of T1D (37,38). We demonstrated that allosteric TYK2is block IFN-mediated transcriptional responses *in vitro* and delay the onset of diabetes in *in vivo* preclinical murine models of T1D. Mechanistic studies, including *in situ* spatial transcriptomics, defined a phenotype whereby pharmacological inhibition of TYK2: 1) dampens innate immune cell activation, 2) impairs adaptive immune responses, 3) directly reduces β cell inflammation and apoptosis, and 4) prevents pathological interactions between T cells and β cells. Taken together, these data support a path forward for testing TYK2is in human T1D.

## RESULTS

### TYK2 inhibitors repress IFNα signaling and inflammatory gene expression in human β cells

To investigate the direct effects of TYK2i treatment on IFNα responses in β cells, islets isolated from human organ donors were treated with IFNα in the presence or absence of BMS-986165 (deucravacitinib), which is FDA-approved for use in psoriasis (33). As anticipated, IFNα treatment stimulated phosphorylation of STAT1 and 2, and this early effect was repressed by the TYK2i in a dose-dependent manner (**Figure 1A-C**). Additionally, co-incubation of human islets with TYK2i (0.3 µM) and combinations of proinflammatory cytokines (IFNɑ+TNFɑ or IFNɑ+IL-1β) prevented the activation of pSTAT1 and pSTAT2 (**Figure 1D-F**). TYK2i also reduced cytokine-induced apoptosis (**Figure 1G**) and decreased the expression of genes involved in ER stress (*DDIT3 (CHOP)* and *ATF3*) (**Figure 1H-I**). Furthermore, the increased expression of a selected signature of IFNα-responsive genes involved in chemokine production (*CXCL10*), antiviral responses (*MX1*), and antigen presentation that contributes to CD8+ T cell target recognition (*HLA-ABC*) was blocked effectively by TYK2i (**Figure 1J-L**). Of note, we have previously shown that TYK2i-induced downregulation of *MX1* does not increase the susceptibility of human β cells to infections by the potentially diabetogenic coxsackievirus B (CVB) (39).

**Figure 1.**
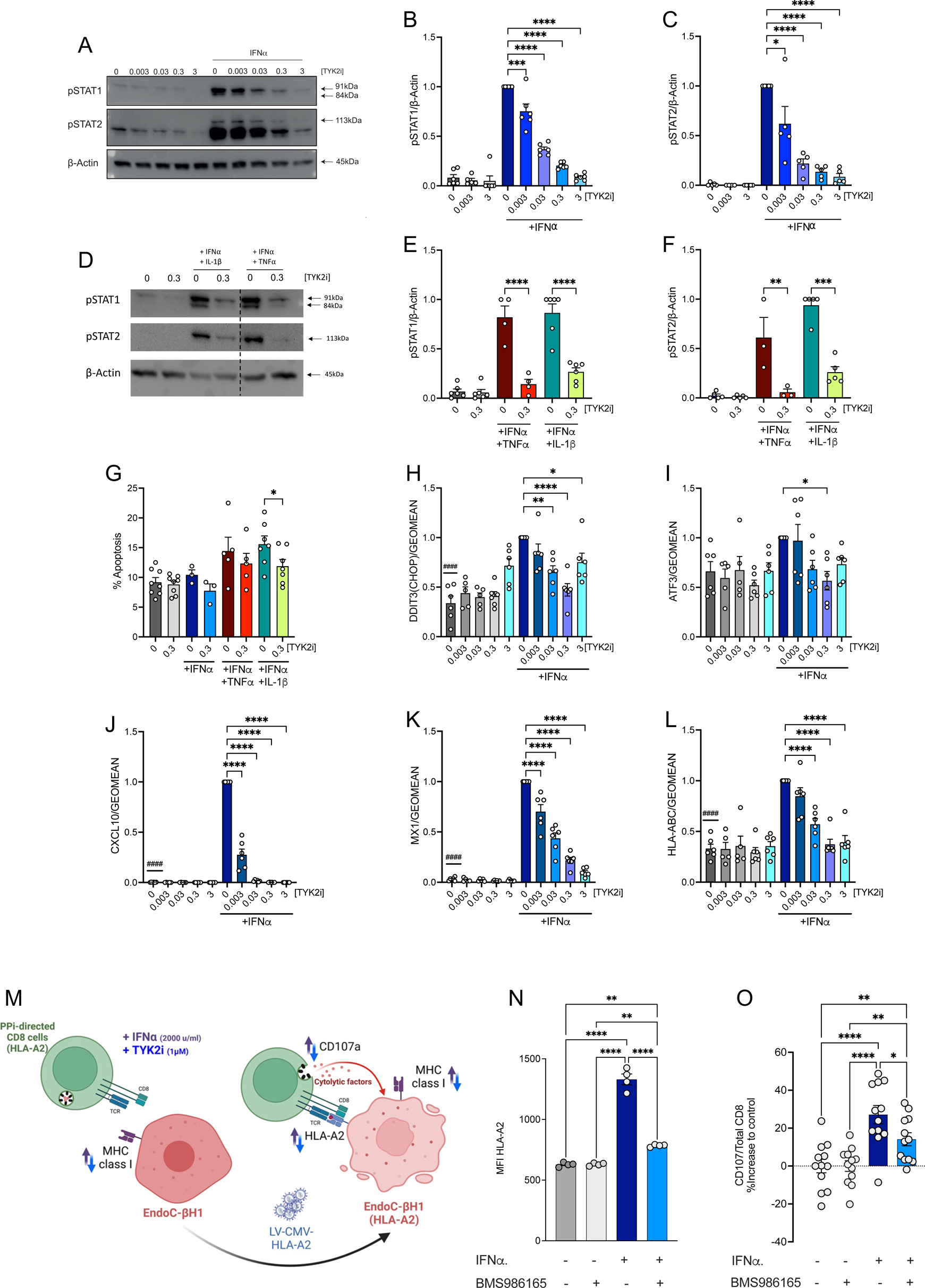
TYK2 inhibitors repress IFNα signaling and inflammatory gene expression and reduce β cell immunogenicity. Dispersed human islet cells were allowed to rest in culture for two days before pre-treatment for 2 h with the indicated concentrations (expressed in mM) of TYK2i BMS-986165. Treatment was continued with BMS-986165 in the absence or presence of IFNα (2000 U/mL), IFNα (2000 U/mL) + TNFα (1000 U/ml), or IFNα (2000 U/mL) + IL-1β (50 U/mL) for 24 h. (**A-F**) Western blotting of protein lysates was performed to detect phospho-STAT1/2. Data were normalized to the levels of β-actin and expressed relative to treatment with IFNα alone. (**G**) Apoptotic cells were identified by Hoechst 33342 and propidium iodide staining. (**H-L**) Total RNA was extracted and mRNA levels of (**H**) DDIT3 (*CHOP*), (**I**) *ATF3*, (**J**) *CXCL10*, (**K**) *MX1*, and (**L**) *HLA-ABC* were analyzed by qPCR. (**M**) Schematic representation of the experimental workflow for measurement of CTL-mediated β cell killing. (**N**) HLA-A2 expression in EndoC-BH1/HLA-A2 upon IFNα treatment (2000 U/mL) in the presence or absence of 1 uM BMS-986165. (**O**) EndoC-βH1/HLA-A2 cocultured with *PPI_15-24_*-specific, HLA-A0201-restricted CD8+ T cells. Prior to coculture, target cells were treated with IFNα in the presence or absence of BMS-986165 for 24 h. T cell activation is shown as relative degranulation, estimated by calculating the percent change of the absolute degranulation of the treated samples compared to their respective controls. For RT-PCR analysis, expression data were normalized by the geometric mean of *GAPDH* and *ACTB* expression levels and expressed relative to cells exposed to IFNα alone. The blotting and expression data are mean ± SEM of 3-7 independent islet preparations per experimental condition; ^####^*p*<0.0001 vs no inhibitor and IFNα (2000 U/mL) and \**p*<0.05, \*\**p*<0.005, \*\*\**p*<0.001, and \*\*\*\**p*<0.0001 vs no inhibitor and with indicated cytokines, one-way ANOVA followed by Bonferroni correction for multiple comparisons. All co-culture experiments were performed with independent biological replicates (n=3); \**p*<0.01; **p<0.007; ****p<0.0001. A one-way ANOVA was used to determine differences between the treatment conditions.

Next, we performed side-by-side comparisons between BMS-986202 and BMS-986165 using preparations of human EndoC-BH1 β cells. The two TYK2is were highly potent, efficacious, and largely similar in their capacity to decrease the expression of IFNα-responsive genes (**Supplemental Figure 1**). TYK2 inhibition also repressed the increase in inflammatory gene expression in response to co-incubation with IFNα+TNFα or IFNα+IL-1β and under conditions of extended IFNα incubation up to 48 hrs (**Supplemental Figure 2**). Notably, the inhibitors did not cause cellular toxicity, as treatment with BMS-986202 or BMS-986165 alone did not increase the percentage of basal apoptosis (**Figure 1G**).

Next, we evaluated the potential for BMS-986202 to modulate the response to IFNα in newly differentiated β cells derived from human inducible pluripotent stem cells (iPSCs). iPSC-derived islet cells are not fully differentiated and provide an interesting model for human islets in the first months of life, a period when β cell autoimmunity may begin in some cases (20). Differentiation of HEL115.6 iPSC cells into endocrine pancreatic cells was performed with quality control evaluations at stages 1, 4, and at end-stage (stage 7), as previously described (40,41). Dispersed islet-like cell aggregates were preincubated in the presence or absence of TYK2i, followed by co-incubation with the inhibitor and IFNα alone or in combination with TNFα or IL-1β for up to 48 hrs. Similar to the results obtained with adult human islet preparations and EndoC-βH1 cells, TYK2i blocked the cytokine-induced increase in the expression of *CXCL10, MX1,* and *HLA-ABC* (**Supplemental Figure 2G-L**), and there was a trend towards decreased apoptosis in iPSCs treated with TYK2i (**Supplemental Figure 2M**).

### TYK2 inhibitors does not alter the expression of β cell identity genes

A recent publication showed that TYK2 knockout in human iPSCs delayed endocrine cell differentiation without impacting mature stem cell-derived islet function (34). Therefore, we tested the impact of BMS-986202 on the expression of β cell identity genes in adult human islets. Importantly, TYK2i treatment did not alter the expression of mRNAs encoding either insulin or PDX1 (**Supplemental Figures 3A-B**).

### TYK2 inhibitors reduce β cell immunogenicity

We have shown that TYK2 inhibition blocks the transcriptional effects of IFNα on β cells; however, it remains to be clarified whether blocking these responses influences interactions, either positive or negative, between β cells and T lymphocytes in the context of insulitis. Among the diverse effects of IFNα, this cytokine induces expression of the checkpoint inhibitor *PD-L1* in β cells. Indeed, we observed that *CD274 (PD-L1)* mRNA expression in human islets was increased by IFNα treatment but repressed by TYK2 inhibition (**Supplementary Figure 3C**). This finding raises the possibility that TYK inhibition could have an untoward effect of heightening deleterious β and T cell interactions, in spite of the parallel decrease in CXCL10 and HLA class I expression induced by TYK2is (**Figures 1J** and **1L**). To test this possibility, we used HLA-A2-specific pre-proinsulin (PPi)-targeting CD8+ T cells to evaluate whether TYK2 inhibition could affect the presentation of the PPi epitope (ALWGPDPAA) to T cells (**Figure 1M**) (42,43). EndoC-βH1 cells expressing HLA-A2 were generated by transduction with a lentiviral vector containing HLA-A02:01 (44). Upon IFNα stimulation, we observed increased HLA-A2 surface expression, as reported previously (45), that was effectively prevented by co-incubation with BMS-986165 (**Figure 1N**). In agreement with these findings, IFNα increased PPi-derived epitope presentation to cytotoxic T lymphocytes (CTLs) and stimulated T cell degranulation, as monitored by CD107a expression at the cell surface, and this effect was reduced by co-treatment with BMS986165 (**Figure 1O**). Therefore, the overall effect of TYK2i on autoimmunogenicity is beneficial, limiting antigenic peptide presentation during inflammation.

Collectively, these *in vitro* results confirm that TYK2i treatment is sufficient to: 1) block the deleterious IFNα-mediated transcriptional effects in isolated human islets, EndoC-βH1 cells, and iPSC-derived β cells without modifying the differentiated β cell phenotype and 2) decrease the susceptibility of β cells to effector T cell attack.

### TYK2i decreases T1D development in the *RIP-LCMV-GP* mouse model

BMS-986202 was selected for *in vivo* studies based on our *in vitro* data (see above) and the compound’s mouse-optimized pharmacokinetic properties (37). First, we tested the impact of BMS-986202 in *RIP-LCMV-GP* mice, a model created by placing the glycoprotein (GP) of the lymphocytic choriomeningitis virus (LCMV) under transcriptional control of the rat insulin promoter (RIP). Upon LCMV inoculation, *RIP-LCMV-GP* mice develop diabetes within 14 days, and disease progression mimics elements of human disease, including dependence upon IFNα signaling and induction of memory CTLs that elicits effector T cell-mediated destruction of β cells (24). In this first series of *in vivo* experiments, 8-week-old *RIP-LCMV-GP* mice were pretreated with vehicle or BMS-986202 (30mg/kg/day) for 2 days and then inoculated with LCMV (0.5 X 10e5 PFU I.P; Armstrong strain) to induce the onset of T1D. Drug or vehicle treatment was continued for 14 days post-LCMV inoculation, and blood glucose levels were measured before LCMV inoculation and on days 1, 4, 7, 11, and 14 post-inoculation (**Figure 2A**). The criterion for established diabetes was two consecutive blood glucose readings exceeding 250 mg/dL.

**Figure 2.**
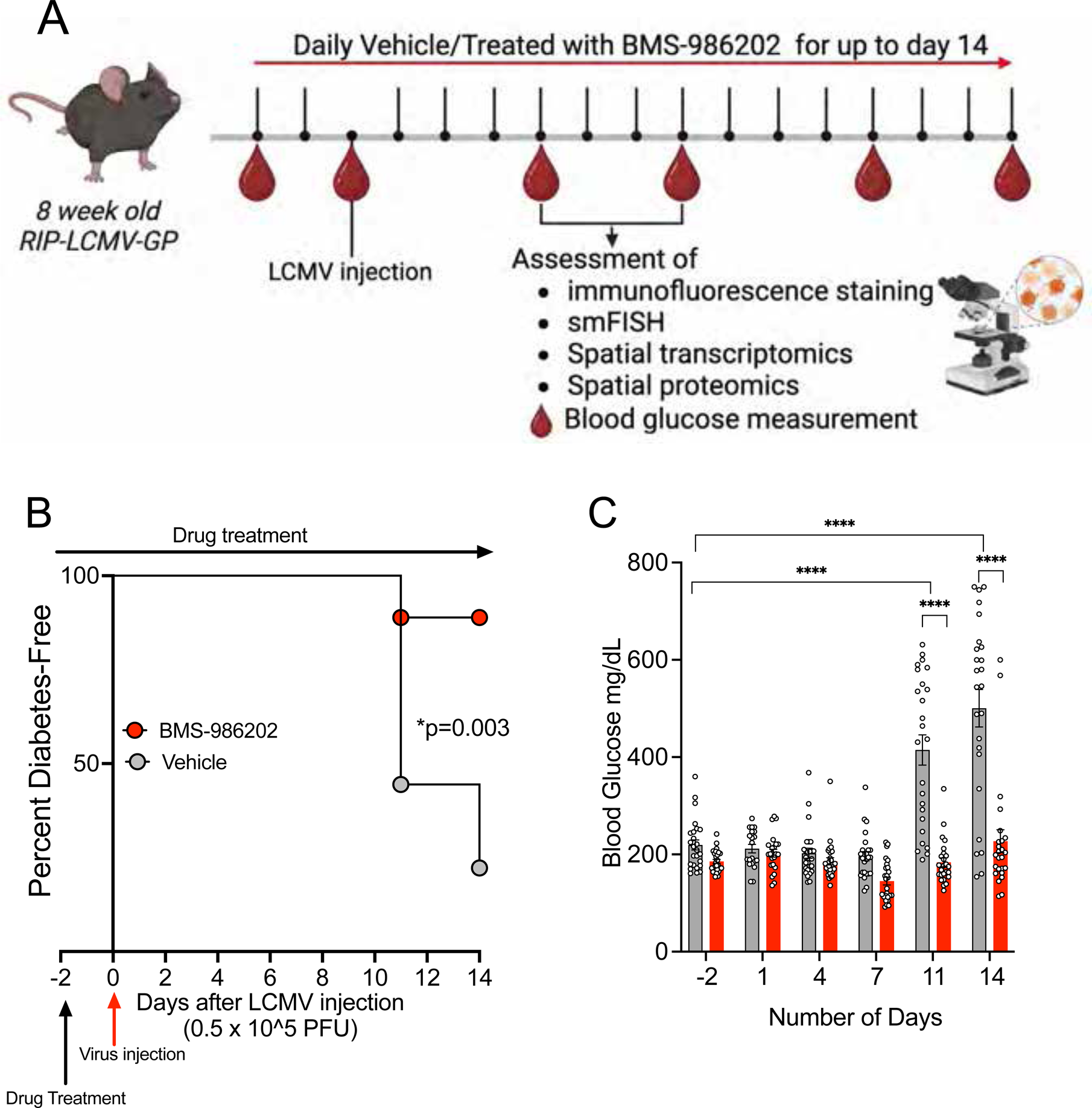
TYK2 inhibition reduces the onset of diabetes in the *RIP-LCMV-GP* mouse model of T1D. (**A**) Male *RIP-LCMV-GP* mice were inoculated at 8 weeks of age with LCMV stock (0.5 X 10^5^ PFU I.P., Armstrong strain), and blood glucose levels were measured pre-inoculation (Pre) and on days 1, 4, 7, 11, and 14 post-inoculation. Mice were pre-treated with either vehicle or TYK2i BMS-986202 (30mg/kg/day) for 2 days prior to inoculation, and treatment continued until the study end on day 14. n=18 mice/group. Vehicle- and TYK2i-treated groups are shown in grey and red symbols or bars, respectively. (**B**) Kaplan-Meier diabetes incidence plot for vehicle- and TYK2i-treated groups. (**C**) Blood glucose levels of vehicle- and TYK2i-treated mice. All blood glucose data are presented as mean ± SEM, and individual data points are included; \**p*=0.003, \*\**p*<0.005, \*\*\**p*<0.001, and \*\*\*\**p*≤0.0001 indicate significant differences determined by Log-rank (Mantel-Cox) test.

Treatment of *RIP-LCMV-GP* mice with BMS-986202 significantly reduced the incidence of diabetes: 89% of BMS-986202-treated mice remained diabetes-free throughout the duration of the study period, while only 14% of the mice in the vehicle-treated group remained diabetes-free (n=18 mice/group, p=0.003; **Figure 2B**). Consistent with the observed protection from diabetes incidence, TYK2i-treated mice had significantly lower blood glucose levels on days 11 and 14 post-inoculation than vehicle-treated mice (**Figure 2C**).

To determine the impact of *in vivo* TYK2 inhibition on β cells, we performed single-molecule fluorescence *in situ* hybridization (smFISH) for a representative set of IFNα-induced mRNAs (*Cd274, Cxcl10, Stat1,* and *Mx1*) and co-stained for insulin protein in pancreatic tissue sections collected from *RIP-LCMV-GP* mice on days 3, 7, and 14 post-LCMV inoculations (**Figure 3**). Consistent with the observed effects on blood glucose, at day 3, there was no difference in insulin protein expression between vehicle- and BMS-986202-treated mice (**Figure 3A**); however, a higher insulin intensity was observed in islets of TYK2i-treated mice on days 7 and 14 (**Figure 3B-C**). Quantification of IFNα-induced mRNAs *in situ* revealed a significant decrease in *Cd274* and *Cxcl10* expression (**Figure 3D-E**). In contrast, we observed an increase in *Mx1* and *Stat1* mRNA expression in TYK2i-treated mice at day 3 post-LCMV inoculation (**Figure 3E**). At day 7, there was a significant decrease in all four mRNAs in β cells of the TYK2i-treated mice (**Figure 3G-H**), and at day 14, a persistent reduction in *Stat1* mRNA was observed in TYK2i-treated mice (**Figure 3J-K**). Analysis of RNA spatial distribution revealed no significant differences in spatial localization (cytoplasmic versus nuclear) of the mRNAs on day 3 (**Figure 3F**) or day 7 (**Figure 3I**). However, at day 14 post-LCMV inoculation, *Cd274* exhibited increased cytoplasmic localization, while *Mx1*, *Cxcl10,* and *Stat1* showed increased nuclear localization in TYK2i-treated mice (**Figure 3L**), suggesting that the spatial distribution of these RNAs might play a role during cellular stress.

**Figure 3.**
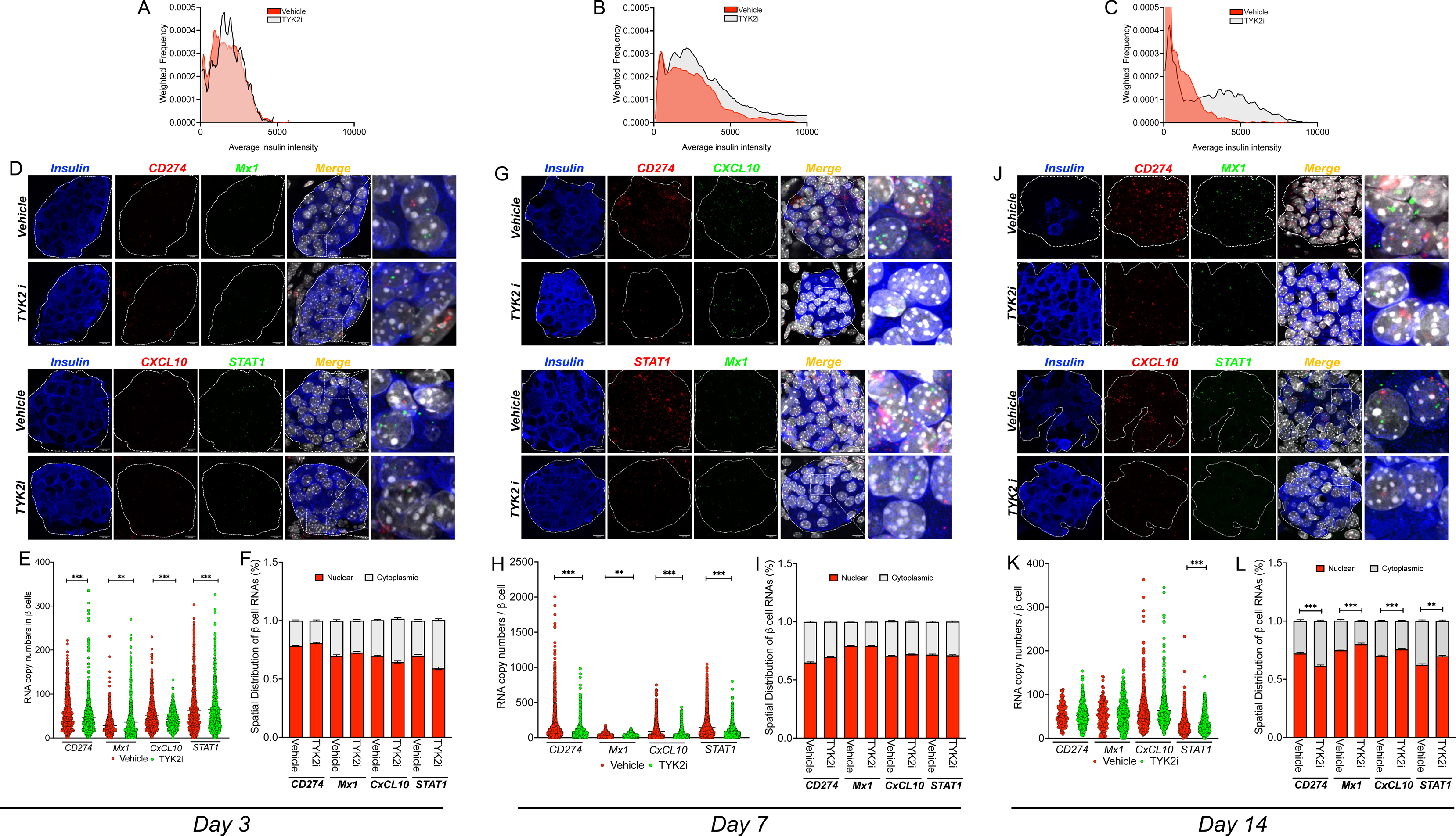
TYK2i treatment preserves pancreatic β cells and inhibits the expression of IFNα-induced mRNAs in *RIP-LCMV-GP* mice. Pancreas tissue was harvested from vehicle- and TYK2i-treated *RIP-LCMV-GP* mice on days 3, 7, and 14 post-inoculation. Single-molecule fluorescence *in situ* hybridization (smFISH) detecting *Cd274*, *Mx1, Cxcl10*, and *Stat1* was performed in combination with co-staining of insulin protein and DAPI for nuclear labeling. mRNA expression in individual insulin-positive β cells was quantified following an established pipeline (details are provided in Methods). (**A-C**) Histogram showing the cellular insulin intensity from the pancreatic tissue sections of vehicle (red plot lines) and TYK2i-treated mice (grey plot lines) on days 3, 7, and 14. **(D-E)** Representative smFISH images of *Cd274*, *Mx1*, *Cxcl10*, and *Stat1* in mouse pancreatic islets on day 3 **(D)**, 7 (**G**), and 14 (**J**). Islet regions are delineated, and the merged smFISH images are shown along with a designated square region of expanded magnification (far right images). (**E, H, K**) Quantitation of mRNA copy number in a single β cell of a pancreatic islet on days 3, 7, and 14. (**F, I, L**) Spatial analysis of RNA localization between nuclear and cytoplasmic subcellular compartments. Individual β cell measurements of RNA copies and RNA localization data are expressed as mean ± SEM with an indication of significant differences, n=5-7 mice were studied per condition and 4-10 islets were randomly selected from each section; \**p*≤0.05, \*\**p*≤0.01, \*\*\**p*≤0.001; statistical significance was determined by Mann-Whitney test.

To understand how these changes in mRNA levels and spatial localization affect protein expression patterns, immunofluorescence staining of PD-L1, CXCL10, and insulin was performed in pancreatic tissue sections at days 3, 7, and 14 post-LCMV inoculation (**Supplemental Figure 3D-E**). Insulin levels in islets from TYK2i-treated mice were transitorily decreased on day 3 compared to vehicle-treated mice (**Supplemental Figure 3E**); however, insulin expression normalized by day 7 and was increased relative to vehicle-treated mice by day 14 (**Supplemental Figure 3E**). Consistent with our smFISH analysis, PD-L1 and CXCL10 protein levels were reduced in β cells of TYK2i-treated mice on both days 3 and 7, indicating early modulation of IFNα induced genes *in vivo* by TYK2i treatment (**Supplemental Figure 3E**).

### TYK2 inhibition decreases early innate immune responses in the whole blood and pancreatic lymph nodes of *RIP-LCMV-GP* mice

To determine if the protective effect of TYK2 inhibition on T1D development in *RIP-LCMV-GP* mice was associated with altered innate or adaptive immune responses, flow cytometry analysis was performed on the whole blood, spleen, and pancreatic lymph nodes (PLN) of vehicle- and TYK2i-treated mice at days 3, 7, and 14 post-LCMV inoculation. The gating strategy for the identification of innate immune cell populations is provided in **Supplemental Figure 4A.** Treatment with TYK2i did not alter the percentage of CD11b^+^ cells in the blood, spleen, or PLN, indicating that the myeloid cell population assayed was largely unaffected (**Figure 4A-C**). However, early in the disease progression (day 3), there was an increase in the percentage of dendritic cells (DCs, CD11b^+^/CD11c^+^/MHCII^+^) in the spleen and a decrease in the percentage of proinflammatory macrophages (F4/80^+^/CD11b^+^/MHCII^+^) in the PLN of TYK2i-treated mice. Furthermore, BMS-986202 treatment decreased the number of mature natural killer (NK) cells (CD49b^+^/CD11b^-^) and increased the subset of immature NK cells (CD49b^+^/CD11b^+^) in the blood (**Figure 4A**). Moreover, at day 7, a critical time point preceding the onset of hyperglycemia in vehicle-treated mice, there was a lower percentage of DCs (CD11c^+^/MHCII^+^) in the PLN of TYK2i-treated mice (**Figure 4B**). These findings indicate that TYK2 inhibition decreases activation of the early innate immune responses in the blood and PLN in this murine model of viral-induced T1D.

**Figure 4.**
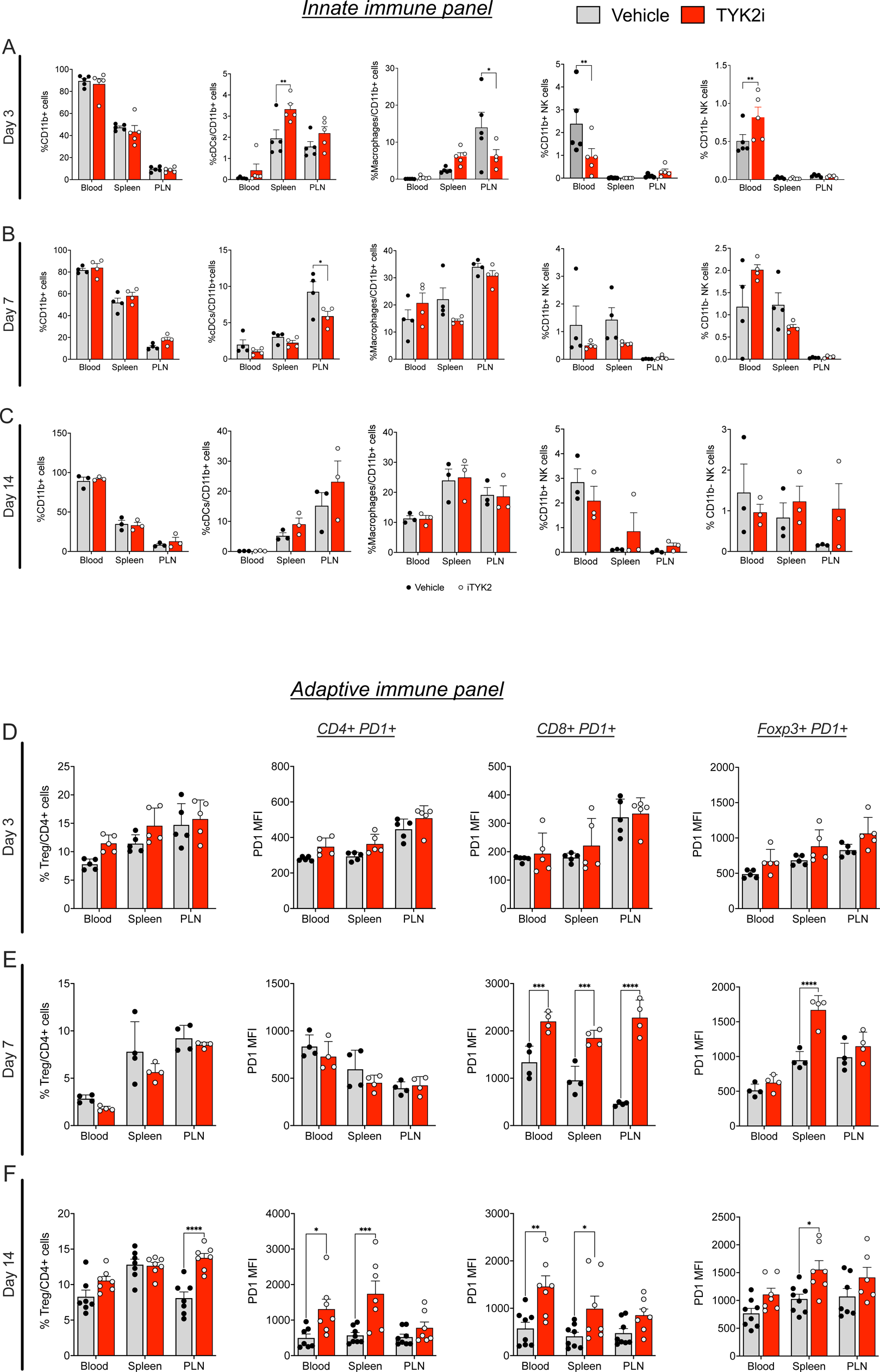
TYK2 inhibition reshapes innate and adaptive immunity in blood, spleen, and PLN RIP-LCMV-GP mice. Peripheral blood mononuclear cells (PBMCs), splenocytes, and immune cells from PLN were isolated and labeled with CD11b^+^MHCII^+^CD11c^+^ (dendritic cells, DCs), CD11b^+^MHCII^+^F4/80^+^ (macrophages), CD11b^+^CD49^+^ (mature NK cells), and CD11b^+^CD49^-^ (immature NK cells). Immune cell characterization was performed using flow cytometry. **(A-C)** Percentage of DCs, macrophages, mature NK cells, and immature NK cells from 3, 7, and 14-days post-inoculation from vehicle (grey) and TYK2i-treated (red) *RIP-LCMV-GP* mice. (**D-F**) Percentage of CD4^+^FoxP3^+^CD25^+^ Tregs, CD4^+^PD1^+^ T cells, and CD8^+^PD1^+^ T cells from 3, 7, and 14 days post-inoculation from vehicle (grey) and TYK2i-treated (red) *RIP-LCMV-GP* mice. Data are presented as mean ± SEM, and individual data points are included, n=3-5 mice per condition; \*\**p*≤0.01, \*\*\**p*≤0.001, \*\*\*\**p*≤0.0001 are indicated for all significant differences. Statistical significance was determined by two-way ANOVA.

### TYK2 inhibition alters the adaptive immune repertoire in the whole blood, spleen, and PLN of *RIP-LCMV-GP* mice

To investigate the impact of TYK2i on adaptive immune responses, we analyzed whole blood, spleen, and PLN on day 3, 7, and 14 post-LCMV inoculation using flow cytometry with the gating strategy described in **Supplemental Figure 4B**. This analysis revealed distinct changes in T cell subsets following TYK2 inhibition but, in contrast to effects on innate immune responses, the effects were observed at the later time points (**Figure 4D-F**). At day 7 post-inoculation, CD8+ T cells from TYK2i-treated mice showed a striking upregulation of PD-1, the receptor of PD-L1, in the blood, spleen, and PLN (**Figure 4E**). This upregulation continued through day 14 in the blood and spleen (**Figure 4F**). At day 14, there was also an increase in the percentage of CD4+PD1^+^ T cells in the blood and spleen of TYK2i-treated mice. Additionally, immune-suppressive regulatory T cells (Tregs, CD25+/FoxP3+) positive for PD-1 (CD25+Foxp3+PD1+) were increased in the spleen of TYK2i-treated mice at days 7 and 14. Moreover, at the 14-day time point, there was a parallel increase in the percentage of Tregs in the PLN (**Figure 4F**). These results suggest that TYK2 inhibition leads to upregulation of PD-1 on CD4+, CD8+, and FoxP3+ cells in a time-dependent manner in *RIP-LCMV-GP* mice.

### TYK2 inhibition attenuates the development of T1D in NOD mice

To investigate the impact of TYK2 inhibition in a spontaneous preclinical model of T1D, female nonobese diabetic (NOD) mice were treated with either vehicle or 30 mg/kg BMS-986202 for 6 weeks beginning at 6 weeks of age. In this model, the spontaneous conversion of female NOD mice to diabetes at approximately 14 weeks of age reflects the onset of insulitis and T cell-mediated destruction of β cells with a high dependence upon IFNα (25,46). Animals treated with vehicle or TYK2i were monitored for blood glucose levels once a week from 6-14 weeks of age and biweekly from 15-24 weeks of age (**Figure 5A**). The criteria for the development of diabetes was two consecutive blood glucose readings that exceeded 250 mg/dL. Pancreatic tissue from vehicle- and BMS-986202-treated mice was collected from a cohort of mice at the end of treatment (12 weeks of age) for histological evaluation of immune cell infiltration into islets. Perivascular/periductal immune cell infiltration and average insulitis score was reduced in the pancreas of mice treated with BMS-986202 compared to vehicle-treated mice (P=0.001, **Figure 5B-C**). By 24 weeks of age, the incidence of diabetes-free mice within the vehicle cohort was less than 25%, whereas 70% of the mice receiving BMS-986202 remained diabetes-free (p<0.0075, **Figure 5D**) with well-maintained glycemia **Figure 5E**.

**Figure 5.**
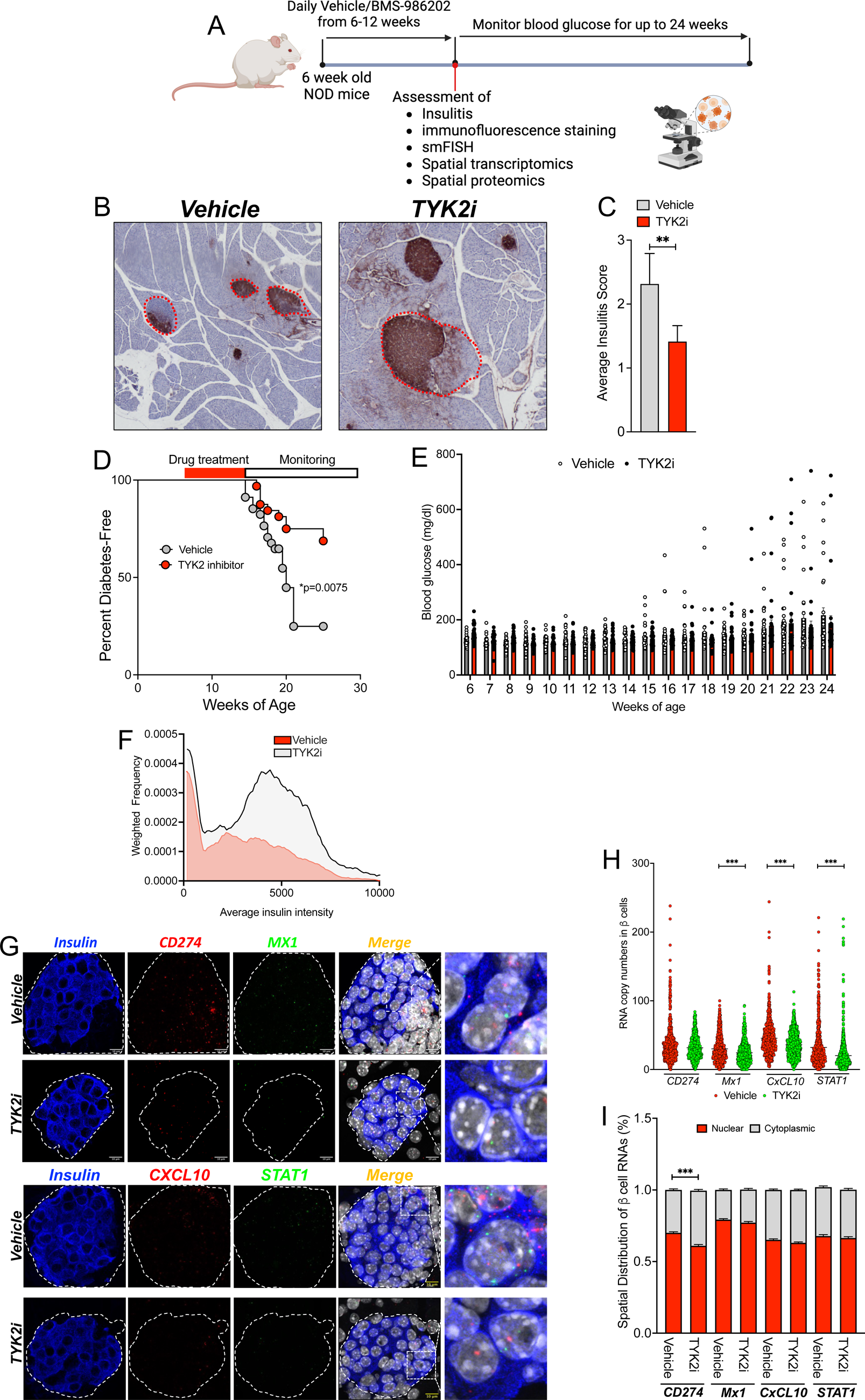
TYK2 inhibition mitigates diabetes onset and IFNɑ responses in female NOD mice. (**A**) Six-week-old female NOD mice were treated with vehicle or TYK2i (BMS986202, 30 mg/kg) by oral gavage once daily for six weeks and monitored for diabetes onset (n=34 vehicle/32 TYK2i). A separate cohort of NOD mice (n=9 per study condition) was sacrificed at the end of TYK2i dosing (12 weeks of age) for tissue analysis. Blood glucose levels were measured weekly from 6 weeks until diabetes conversion (14 weeks) and biweekly until the end of the study time point. Diabetes was defined as blood glucose levels of ≥250 mg/dL in two consecutive measurements. (**B-C**) Representative images of insulin immunohistochemistry and insulitis scoring. (**D**) Kaplan-Meier diabetes incidence plot. (**E**) Blood glucose levels. Vehicle-treated mice are indicated in grey bars and TYK2i-treated mice are represented by red bars. (**F**) Quantification of insulin expression in pancreatic β cells. (**G**) Representative images of smFISH for *Cd274*, *Cxcl10*, *Stat1*, and *Mx1*. Islet regions are delineated, and the merged smFISH images are included along with a designated square region of expanded magnification (far right images). (**H**) Quantitation of mRNA copy numbers of *Cd274*, *Cxcl10*, *Stat1*, and *Mx1 in* β cells. (**I**) Spatial analysis of RNA localization between nuclear and cytoplasmic cellular compartments. For diabetes incidence studies, statistical analysis was performed by Log-rank (Mantel-Cox) test for Kaplan-Meier plot and one sample t-test and Wilcoxon test. For smFISH, individual β cell measurements of RNA copy are presented, RNA localization data are expressed as mean ± SEM, and the Mann-Whitney test was used to determine statistical differences; *p=0.0075; **p<0.001; \*\*\**p*≤0.0001.

Similar to the analysis performed in the *RIP-LCMV-GP* mice, smFISH was performed on the pancreas from 12-week-old vehicle- and TYK2i-treated NOD mice to detect IFNα-mediated inflammatory gene expression in β cells (**Figure 5F-I**). Overall, the signature of inflammatory gene expression was reduced in β cells from TYK2i-treated mice, with significantly lower levels of *Mx-1*, *CxcL-10*, and *Stat1* mRNAs (**Figure 5G**), and the cytoplasmic localization of *Cd274* mRNA was significantly increased (**Figure 5H**). Immunofluorescence staining of pancreatic tissue sections for PD-L1, CXCL10, and insulin did not reveal significant differences between the groups at 12 weeks of age (**Figure 5F** and **Supplemental Figure 5A-B**).

### Evaluation of spatial dynamics reveals that TYK2 inhibition reshapes inflammatory signaling and cellular composition in T1D models

We reasoned that an in-depth examination of the impact of TYK2 inhibition across target tissues would provide additional insight into their effects on disease pathogenesis. Therefore, we used a GeoMx spatial whole transcriptomic atlas (WTA) to evaluate islets, spleen, and PLN from *RIP-LCMV-GP* mice at days 3 and 7 post-inoculation (**Figure 6A-S** and **Supplemental Figure 6**) and islets and PLN from NOD mice treated with vehicle or TYK2i (**Figure 7A-J** and **Supplemental Figure 7**). GeoMx WTA probes were hybridized, and the tissue sections were stained for insulin, CD3, and CD68 to select regions of interest (ROIs) (**Figure 6A-B** and **Figure 7 A-B**). A total of 193 ROIs were selected from *RIP-LCMV-GP* mice, and 92 ROIs were selected from NOD mice. After filtering for probe and segment QC, we retained 182 ROIs from *RIP-LCMV-GP* mice and 92 ROIs from NOD mice, on which 5268 genes in *RIP-LCMV-GP* mice (**Supplemental Figure 6**) and 19873 genes in NOD mice (**Supplemental Figure 7**) were quantified. In both *RIP-LCMV-GP* mice and NOD mice, all the ROIs were clustered by tissue and treatment status (**Figure 6C** and **Figure 7C**).

**Figure 6.**
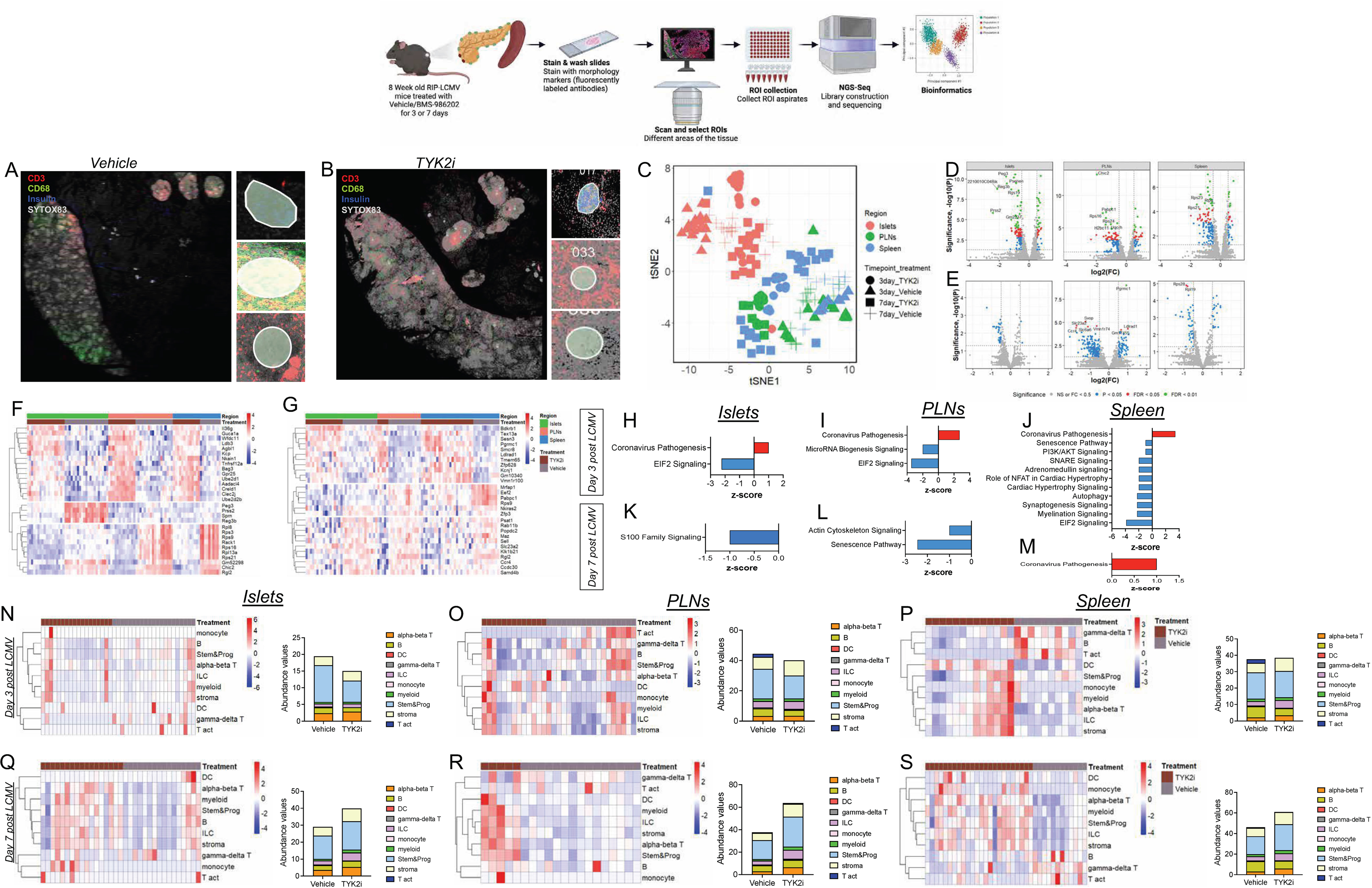
TYK2 inhibition alters spatial transcriptome profiles in islets, PLN, and spleen of *RIP-LCMV-GP* mice. Tissues were harvested from vehicle- and TYK2i-treated *RIP-LCMV-GP* mice on days 3 and 7 post-inoculation. (**A-B**) Representative images of the islets, PLN, and spleen stained for CD3 (red), CD68 (green), insulin (blue), and Sytox83 (grey) nuclear staining. (**C**) tSNE scatter plot of sample clustering between vehicle- and TYK2i-treated mice at days 3 and 7 post-inoculation in the islets, PLN, and spleen. (**D-E**) Volcano plots of differentially expressed genes between vehicle- and TYK2i-treated mice at (**D**) day 3 and (**E**) day 7 post-inoculation. (**F-G**) Top 10 differentially expressed genes from each selected ROI of vehicle- and TYK2i-treated mice at (**F**) day 3 and (**G**) day 7 post-inoculation. (**H-M**) Ingenuity pathway analysis showing upregulated (red bar) and downregulated (blue bar) pathways in islets, PLN, and spleen at (**H-J**) day 3 and (**K-M**) day 7 post-inoculation. (**N-S**) Deconvolution analysis of immune cells showing the abundance of different immune cell populations from islets, PLN, and spleen at (**N-P**) day 3 and (**Q-S**) day 7 post-inoculation. n=3-4 mice per group.

**Figure 7.**
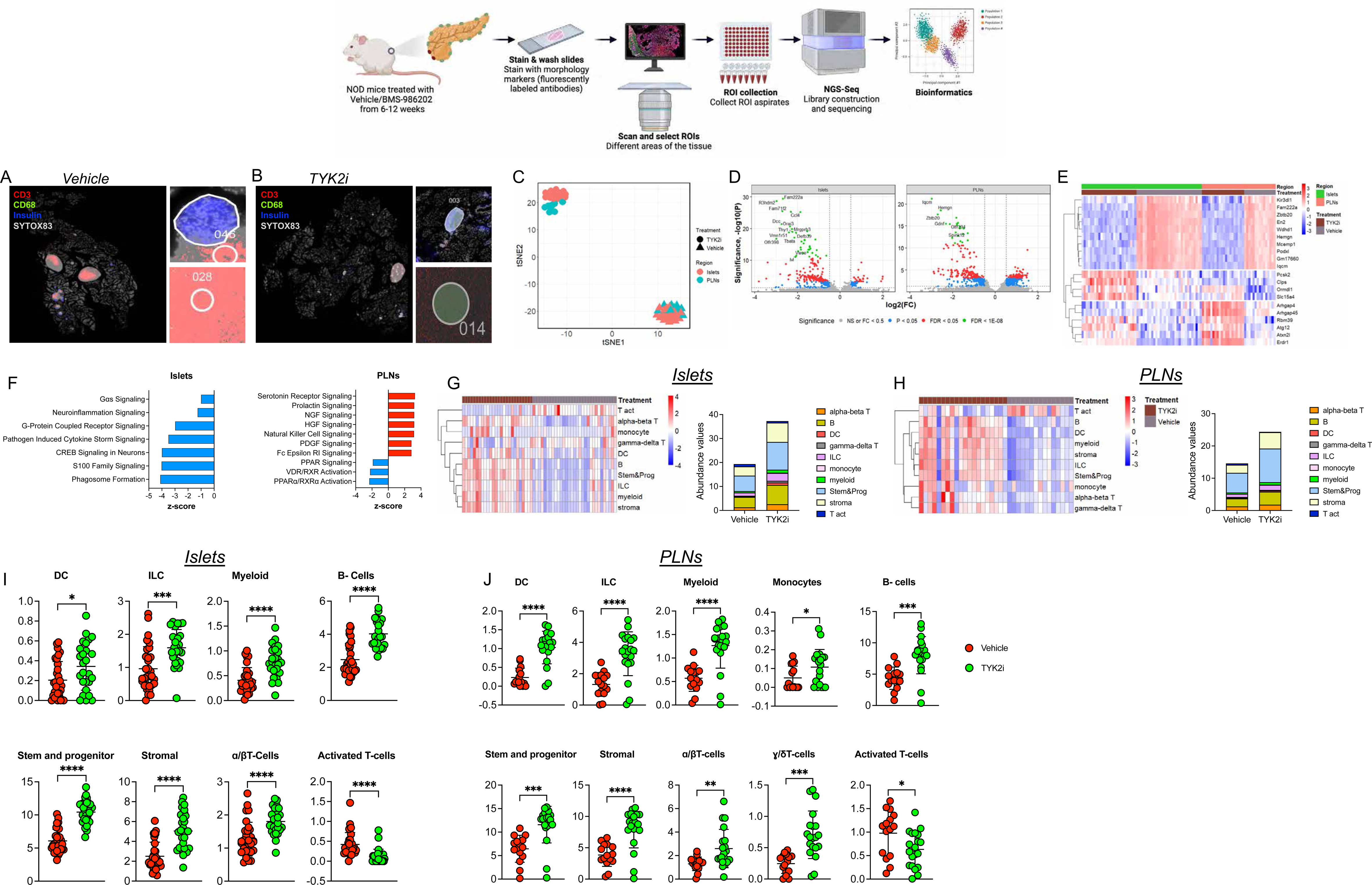
Spatial whole transcriptome analysis reveals diminished inflammatory gene expression in TYK2i-treated islets and PLN of NOD mice. Islets and PLNs were harvested from vehicle- and TYK2i-treated NOD mice at 12 weeks of age. (**A-B**) Representative images of islets and PLN stained for CD3 (red), CD68 (green), insulin (blue), and Sytox83 (grey; nuclear staining). (**C**) tSNE scatter plot showing clustering of samples from islets and PLN between vehicle- and TYK2i-treated 12-week-old female NOD mice. (**D**) Volcano plots showing differentially expressed genes from islets and PLN between vehicle- and TYK2i-treated mice. (**E**) Top 10 differentially expressed genes from each selected ROI in islets and PLN of mice treated with vehicle or TYK2i. (**F**) Ingenuity pathway analysis showing upregulated (red bars) and downregulated (blue bars) pathways in islets and PLN of NOD mice. (**G-J**) Deconvolution analysis of immune cells showing the abundance of different immune cell populations from (**G**) islets and (**H**) PLN from NOD mice treated with vehicle or TYK2i; abundance of immune cell populations showing significant differences between vehicle- and TYK2i-treated NOD mice from (**I**) islets and (**J**) PLN. n=3-4 mice/group. Dotplot showing the proportion of cell types present in each ROI identified by spatial deconvolution. Unpaired t-test was used to compare the statistical significance between vehicle- and TYK2i-treated groups; *p<0.01, **p<0.001, ***p<0.0005 and ****p<0.0001.

Next, we investigated gene expression profiles within islets, spleen, and PLNs following TYK2 inhibition. At day 3 in *RIP-LCMV-GP* mice, we identified 102, 118, and 117 differentially expressed genes from islets, PLN, and spleen, respectively, compared to 16, 201, and 39 differentially expressed genes identified in islets, PLN, and spleen at day 7. Notably, in *RIP-LCMV-GP* mice at day 3 post-LCMV administration, there was a significant upregulation of genes associated with autophagy (*Bag3*) and anti-inflammatory genes (including *Kcp*) in islets, PLN, and spleen, suggesting a potential enhancement in cellular self-regulation and attenuation of inflammation in response to TYK2 inhibition (**Figure 6F**). In contrast, expression of genes linked with tissue damage (*Reg3b*), pancreatitis (*Prss2*), and p53-mediated apoptosis (*Peg3*) were decreased in TYK2i-treated mice (**Figure 6F**), suggesting activation of protective mechanisms. Moreover, an upregulation of several ribosomal genes (*Rack1*, *Rpl8*, *Rps3*, *Rps9*, *Rps16*, *Rpl13a*, and *Rps21*) in the spleen and PLN of the TYK2i-treated group was observed (**Figure 6F**), suggesting that TYK2 inhibition may impact stress-mediated translational responses.

At day 7 post-LCMV inoculation, the differences in gene expression profiles persisted, indicating a lasting impact of TYK2 inhibition (**Figure 6G**). In islets, PLN, and spleen, TYK2i-treated mice exhibited a notable decrease in the expression of sterile alpha motif (SAM) (*Samd4b*), which is involved in the regulation of the transcriptional activity of p53, p21, and chemokine receptor 4 (47). Importantly, there was downregulation of *Eef2*, which is linked to T cell exhaustion, in the PLN and spleen of TYK2i-treated mice. This finding was in line with our observation of increased CD4^+^PD-1^+^ and CD8^+^PD-1^+^ T cells in the flow cytometry analysis of these tissues. Interestingly, TYK2 inhibitor treatment decreased the expression of RNA binding protein (*Pabpc1*) (48,49) in PLN and spleen (**Figure 6G**), suggesting decreased ER stress and further emphasizing the beneficial role of TYK2 inhibition in immune and stress regulation.

Pathway analysis of differentially expressed genes from the islets, PLN, and spleen at days 3 and 7 post-LCMV inoculation suggested that TYK2 inhibition prevented the activation of major inflammatory pathways such as EIF2 signaling, microRNA biogenesis signaling, and senescence signaling (**Figure 6H-M**). There was an activation of coronavirus pathways in all tissues of the TYK2i-treated mice, potentially indicating a lower efficiency in LCMV viral clearance. We next extended our investigation to NOD mice (**Figure 7**). In this model, TYK2i treatment resulted in the downregulation of genes associated with innate immune cell proliferation (*Hemgn*, *Mcemp1*, and *En2*)(50–54). In addition, there was increased expression of genes associated with the negative regulation of the Rho signaling pathway (*Arhgap4* and *Arhgap45*) (55,56), which has been shown to modulate immune cell viability, glucose uptake, and lactate release (**Figure 7D**). Similar to our findings in the *RIP-LCMV-GP* model, we observed upregulation of autophagy-related genes (*Atg12*) and genes involved in the negative regulation of inflammation (*Erdr1*) in islets and PLN of TYK2i-treated mice (**Figure 7E**).

To gain insight into the functional implications of TYK2 inhibition in NOD mice, we performed an Ingenuity Pathway Analysis (**Figure 7F**). Within islets, TYK2i treatment led to the downregulation of major inflammatory pathways, including neuroinflammatory signaling, pathogen-induced cytokine storm signaling, S100 family signaling, CREB signaling, and Gɑs signaling. Conversely, our analysis of PLN revealed the downregulation of pathways regulating apoptosis and DNA damage in immune cells (RXR signaling and PARP signaling). Most intriguingly, TYK2i treatment led to the upregulation of NK cell signaling and NGF and HGF signaling pathways in PLN (**Figure 7F**). These findings suggest that inhibition of TYK2 modulates immune responses within lymph nodes in NOD mice.

### Spatial deconvolution analysis shows that TYK2 inhibition modulates immune cell populations in T1D models

Next, we employed spatial deconvolution analysis to determine the evolving dynamics of immune regulation in response to TYK2 inhibition in islets, PLNs, and spleen at days 3 and 7 during the evolution of LCMV-induced T1D in the *RIP-LCMV-GP* mice (**Figure 6N-S** and **Supplemental Figure 6E-J**). On day 3 post-LCMV injection, we observed a significant reduction in gamma delta (ɣδ) T cells and a decline in stem and progenitor cells (**Supplemental Figure 6E**). Simultaneously, a shift in immune cell populations was seen in the spleen and PLNs, where there was a substantial decrease in B cell populations and an increase in DCs in PLN and spleen of TYK2i-treated mice. However, there was a significant decrease in activated T cells in PLN at day 3 in TYK2i-treated mice. Furthermore, TYK2i treatment increased ɑβ T cells in PLN and spleen and induced a significant decrease in ɣδ T cells in the spleen at day 3, suggesting that TYK2 inhibition results in complex immune cell modulation in multiple target tissues (**Supplemental Figure 6I**).

On day 7 post-LCMV injection, we observed a substantial increase in ɑβ T cells, innate lymphoid cells (ILCs), monocytes, myeloid cells, and stromal cells within islets of TYK2i-treated mice. In the spleen and PLN, there were profound alterations in immune cell profiles, with an increase in DC, ILC, myeloid cell, stem and progenitor cell, and stromal cell populations. Consistent with the day 3 effects, there was a significant decrease in B cell and ɣδ T cell populations in the spleen of TKY2i-treated mice on day 7 (**Supplemental Figure 6J**).

In agreement with the above observations in the *RIP-LCMV-GP* model, data from NOD mice corroborated further the effects of TYK2i treatment on the tissue immune cell repertoire (**Figure 7I-J** and **Supplemental Figure 7**). A significant decrease in activated T cells was noted in islets and PLN of TYK2i-treated mice, accompanied by an increase in DCs, ILCs, myeloid cells, stem and progenitor cells, stromal cells, ɑβ T cells, and ɣδT cells (**Figure 7I-J**). This consistency in immune cell population dynamics across two mouse models of T1D underscores the robustness of the effects induced by TYK2 inhibition across the pancreas, PLN, and spleen.

### Spatial proteomics shows that TYK2 inhibition reshapes immune dynamics in T1D models

To substantiate further our findings, we performed GeoMx spatial proteomics profiling on islets, PLNs, and spleen collected from *RIP-LCMV-GP* mice at 3 and 7 days post-LCMV administration (**Figure 8A-H** and **Supplemental Figure 8**) and on islets and PLN from NOD mice at 12 weeks of age (**Figure 8I-N**). Tissue sections from 3-4 mice per group were stained with CD3, B220, insulin, and Sytox83 (nucleus) to identify ROIs. At day 3 post-LCMV infection, no discernible differences were observed in the proportion of immune cells or markers of immune cell activation between the vehicle- and TYK2i-treated *RIP-LCMV-GP* mice (**Supplemental Figures 8**). However, by day 7 post-LCMV infection, in TYK2i-treated mice, there was a significant increase in the proportion of cells positive for CD11b and CD11c in PLN, and there was also an increase in the number of cells expressing Ki67, CD127/IL7RA, and CD40L in PLN and spleen compared to vehicle-treated mice. Notably, a heightened presence of FoxP3-positive cells was noted in the spleen of TYK2i-treated mice. We also observed an upregulation of PD-1 expression in the PLN and spleen of TYK2i-treated mice (**Figure 8F-H**). These findings are consistent with our flow cytometry and spatial deconvolution data and collectively suggest that TYK2 inhibition in *RIP-LCMV-GP* mice curbs T cell activation while leaving innate immune cell profiles largely unaffected.

**Figure 8.**
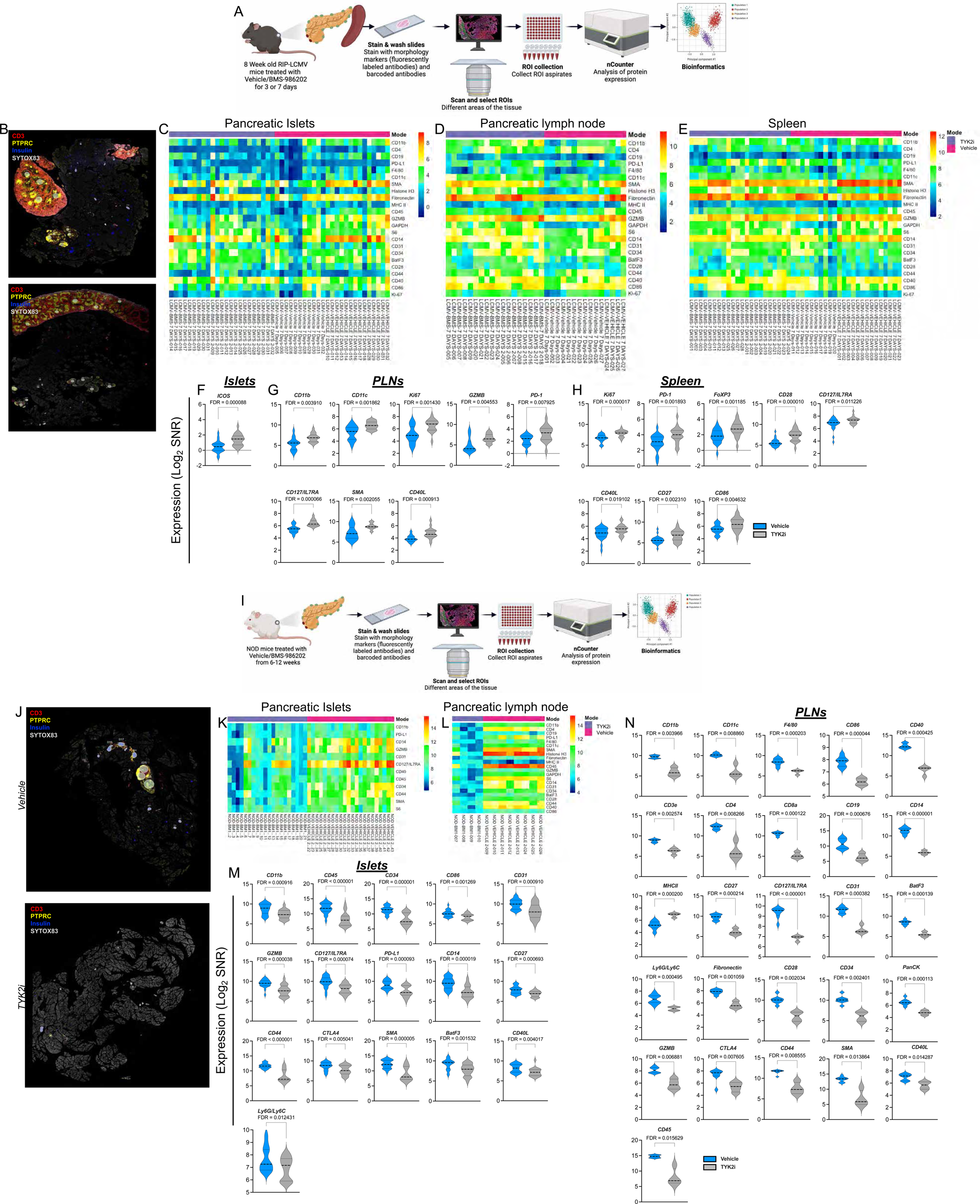
Spatial proteomics uncovers immune cell dynamics in TYK2i-treated *RIP-LCMV-GP* and NOD mouse models. Tissues were harvested from vehicle- and TYK2i-treated *RIP-LCMV-GP* mice on day 7 post-inoculation. (**A**) Schematic representation of GeoMx-DSP proteomics experimental workflow from RIP-LCMV-GP mice. (**B**) Representative images of islets, PLN, and spleen labeled for CD3 (red), PTPRC (yellow), insulin (blue), and Sytox83 (grey). (**C-E**) Heatmap showing the overall expression of immune cell typing and immune cell activation makers from selected ROIs in (**C**) islets, (**D**) PLN, and (**E**) spleen. (**F-H**) Markers that show a significant difference between vehicle- and TYK2i-treated mice in (**F**) islets, (**G**) PLN, and (**H**) spleen. (**I**) Schematic representation of GeoMx-DSP proteomics experimental workflow for tissues from NOD mice. (**J**) Representative images of islets and PLN labeled for CD3 (red), PTPRC (yellow), insulin (blue), and Sytox83 (grey). (**K-L**) Heatmap showing the overall expression of immune cell typing and immune cell activation markers from selected ROIs in (**K**) islets and (**L**) PLN. (**M-N**) Markers that showed significant differences between vehicle- and TYK2i-treated mice from (**M**) islets and (**N**) PLN; n=4 mice/group for RIP-LCMV-GP mice and n=3-4 for NOD mice per group. Data were normalized to the geometric mean of the IgG negative control, and the significantly differently expressed proteins were presented as Log_2_ of signal-to-noise ratio (SNR) and the violin plots were presented with FDR.

In NOD mice treated with TYK2i there was a substantial reduction in markers of immune cell activation and immune cell populations in islets and PLN (**Figure 8I-N**), with a significant decrease in the expression of innate and adaptive immune cell markers (CD11b, CD11c, F4/80, CD86, CD40, CD3e, CD4, CD8a, CD19, CD14, CD27, CD127/IL7RA, CD31). In addition, we also observed a significant decrease in the expression of markers of T cell activation (Ly6G/Ly6C, Fibronectin, CD28, CD34, PanCK, GZMB, CTLA4, CD44, SMA, CD40L, and CD45) in TYK2i treated NOD mice. Despite the decreased expression of activation markers, an increase in MHCII expression was observed in PLN of NOD mice receiving TYK2i treatment (**Figure 8N**).

## DISCUSSION

The first small molecule TYK2 inhibitor was approved by the FDA in 2022 for the treatment of psoriasis, with additional agents under investigation for use in other autoimmune conditions (57–59). TYK2 mediates signaling through several cytokines that have been linked with T1D pathogenesis, including IFNα, IL-12, and IL-23. Along these lines, missense mutations in TYK2 are associated with protection against T1D development, making it an appealing pharmacological target for disease intervention (3,6). In the present study, we demonstrate that the highly specific TYK2 pseudokinase inhibitors, BMS-986165 and BMS-986202, modulate three critical nodes in T1D pathophysiology including: 1) immune cell activation and target tissue infiltration; 2) β cell inflammation and survival; and 3) direct interaction of β cells with antigen-specific CD8+ T cells. Taken together, these data provide a robust rationale for testing the effectiveness of TYK2 inhibitors in clinical trials to prevent and/or delaying the development of T1D.

We utilized three *in vitro* human experimental models (EndoC-βH1 cells, dispersed adult human islets, and pancreatic islet-like cells differentiated from human iPSCs) to study the direct effects of TYK inhibition on the β cell. We found that BMS-986165 and BMS-986202 reduced IFNα-mediated upregulation of *CHOP*, *HLA class I, MX1*, and *CXCL10* gene expression. Similarly, TYK2 inhibition reduced β cell chemokine production, ER stress, and HLA Class I upregulation in response to a cocktail of cytokines that included IFNɑ+TNFɑ and IFNɑ+IL-1β. These data are consistent with a previous publication from our group using two different TYK2is (34,39) and findings from others using BMS-986165 (60). Importantly, we show these effects in the β cell are sufficient to reduce T cell-mediated β cell killing in a co-culture system.

To test the *in vivo* efficacy of TYK2i, we utilized two well-established mouse models of T1D. We found that TYK2 inhibition prevents the onset of hyperglycemia in both RIP-*LCMV-GP* and NOD mice. We leveraged the predictable and rapid onset of T1D with viral induction in the *RIP-LCMV-GP* model to define the temporal effects of TYK2 inhibition. At early time points (i.e., 3 days after viral induction), flow cytometry revealed a decrease in mature NK cells in the blood of TYK2i-treated *RIP-LCMV-GP* mice, with a reciprocal increase in a subset of immature NK cells. This particular finding was notable given previous studies showing changes in NK cell phenotypes in established T1D (61) and the recent identification of an NK-enriched signature associated with both islet autoimmunity and T1D onset in the TEDDY and DIPP cohorts, which follow newborns with high genetic risk of T1D (62,63). In addition, we observed an increase in the percentage of CD11b+ DCs in the spleen of TYK2i-treated *RIP-LCMV-GP* mice, followed by a reduction of these cells in the PLN at day 7. Intriguingly, spatial transcriptomics indicated an increase in DC populations that expressed CD207 in the islets and PLN of TYK2i-treated mice. While the exact phenotypes and roles of DCs in T1D have been controversial, the expansion of specific DC populations, including those that express CD207, has been linked with pro-tolerogenic phenotypes (64,65).

At later timepoints, immune profiling of TYK2i-treated *RIP-LCMV-GP* mice revealed changes that were largely restricted to adaptive immune cell subsets. In this model, flow cytometry revealed increased percentages of PD1^+^ CD4^+^ and CD8^+^ T cells in the blood, spleen, and PLN of TYK2i-treated mice, while spatial transcriptomics showed decreased ɣ/δT-cells in the islets, spleen, and PLN. Consistent with changes in effector cell subsets, transcriptomics in TYK2i-treated NOD mice revealed decreased expression of markers of immune cell activation (CD86, GZMB, CD127/IL7RA, CD44, CTLA4, SMA, BatF3, CD40L, and Ly6G/Ly6C) in both islets and PLN, coupled with reduced insulitis. The effects of TYK2i on PD1-expressing T cell subsets were striking, as chronic activation of PD1 is linked with T cell exhaustion characterized by reduced T cell activation (66), diminished cytokine production, and reduced cytotoxic activity (67). Importantly, the most effective immune modulator for T1D prevention to date, Teplizumab, is also associated with an exhausted T cell phenotype (68), and T cell exhaustion has also been linked with response to anti-thymocyte globulin (ATG) treatment in Stage 3 T1D (69,70). Intriguingly, we observed an increase in FoxP3^+^PD1^+^ Tregs in the spleen and PLN in TYK2i-treated *RIP-LCMV-GP* mice. Alterations in Treg function are a well-described component of T1D pathophysiology (71); however, relatively few studies have characterized the frequency or function of PD1^+^ expressing Tregs in models of diabetes. NOD mice with inducible deletion of PD1^+^ were protected against diabetes (72), suggesting that PD1 may restrain Treg function in models of autoimmunity. In contrast, in cancer models, PD1-expressing Tregs possess potent immunosuppressive effects and are associated with resistance to checkpoint inhibitors. In human cohort studies, the ratio between PD1^+^ effector and regulatory subsets has been proposed as a biomarker of cancer drug response (63), indicating there is complexity in interactions across different immune cell subsets that may not be appreciated when PD1^+^ is deleted in a single immune cell type.

At the level of the pancreas, *in vivo* TYK2i treatment modulated expression patterns of interferon-stimulated (ISG) genes, with reduced *CD274, MX1, CXCL10*, and *STAT1* in *RIP-LCMV-GP* mice and reduced *Mx1, CXCL10*, and *STAT1* in NOD mice. TYK2i treatment was associated with changes in the subcellular localization of key ISGs. Most notably, cytoplasmic CD274 mRNA was increased in TYK2i-treated *RIP-LCMV-GP* mice at day 14 and in NOD mice, suggesting that changes in subcellular localization of mRNAs may be influenced by reducing inflammation (74). Of note, changes in mRNA localization have been linked with changes in mRNA translation in several disease models (75,76). Interestingly, PDL-1 protein was reduced in islets from TYK2i-treated *RIP-LCMV-GP* mice at day 3 and 7, but levels were not different between vehicle- and TYK2i-treated mice at day 14. Similar expression patterns of PDL-1 were observed in NOD mice.

Spatial transcriptomics allowed us to compare tissue-specific signatures across the islets, PLNs, and spleen and probe novel pathways through which TYK2is may impact the islet. The number of differentially expressed genes was similar across these tissues, highlighting the potential of this class of drugs to modulate signaling in both the immune and endocrine compartments. Pathway analysis highlighted downregulation of pathways related to EIF2A signaling, S100A signaling, and senescence in TKY2i-treated *RIP-LCMV-GP* mice, while inflammatory pathways, including CREB and Gαs signaling, were downregulated in NOD mice treated with TYK2i.

In a recent publication (36), Mine et al studied the phenotype of NOD mice with total body TYK2 deletion and found that these mice were protected against T1D development and exhibited decreased activation of autoreactive CD8^+^ T-BET^+^ CTLs due to reduced IL-12 signaling in CD8^+^ T cells. Additionally, in a limited pharmacological prevention study, the authors found that NOD mice treated with TYK2i BMS-986165 from 6-10 weeks of age were partially protected from diabetes development. Importantly, the major findings of this study are in agreement with our observations, underscoring the reproducibility and clinical potential of modulating this pathway. However, our study also characterizes how TYK2is impacts β cell stress pathways and how they modulate innate and adaptative immune responses at multiple levels, from the transcriptome to the proteome, with detailed longitudinal phenotyping, thus adding substantially to our understanding of the role of this pathway in the development of T1D. Furthermore, our validation in three human β cell models and two mouse models support the translation of these agents for clinical testing in polygenic and heterogenous populations.

We acknowledge that several key questions remain as to the ideal timing and chronicity of intervention with TYK2 inhibitors. For example, should testing of TYK2is be prioritized in at-risk individuals (i.e. Stage 1 or 2) or after Stage 3/clinical disease onset? The fact that an IFN signature is present in β cells in both the pre-diabetic period and after disease onset (10,62) suggests that treatment at earlier timepoints could be more efficacious to preserve β cell mass and function. However, the recent report that a JAK inhibitor was able to preserve C-peptide levels in Stage 3 T1D (77) provides support for initial testing of TYK2i after clinical disease onset. Second, the duration of therapy is an important consideration for cytokine inhibitors. Data from our preclinical studies suggest that there is modulation of the adaptive immune cell repertoire, but whether this is a persistent effect remains to be tested. The application of JAK and TYK2 inhibitors in other autoimmune conditions suggests chronic dosing would likely be necessary (78,79). Notwithstanding these unresolved questions, our results underscore the diverse and beneficial effects of TYK2 inhibition at the level of the β cell and the immune system in pre-clinical models of T1D, strongly supporting the rationale for testing these agents as a novel therapy to preserve β cells in T1D.

## Supporting information

Human Islet Checklist

STAR Methods

## Acknowledgements and Funding

This work was supported by NIH grants R01 DK093954 and DK127308 (to CEM) and U01DK127786 and UC4DK104166 (to CEM and DLE), VA Merit Award I01BX001733 (to CEM), 2-SRA-2019-834-S-B, JDRF 2-SRA-2018-493-A-B, 3-IND-2022-1235-I-X (to CEM and DLE), and gifts from the Sigma Beta Sorority, the Ball Brothers Foundation, and the George and Frances Ball Foundation (to CEM). FS was supported by JDRF postdoctoral fellowship (3-PDF-20016-199-A-N and JDRF 5-CDA-2022-1176-A-N). This work is also supported by NIH/NIDDK dkNET U24DK097771 (to FS and JL). SAW was supported by an F31 Fellowship from the National Institute of Diabetes and Digestive and Kidney Diseases (NIDDK) award F31DK134168 and a TL1 fellowship from the National Center for Advancing Translational Sciences, Clinical and Translational Sciences Award UL1TR002529. The authors acknowledge the support of the Islet and Physiology Core and the Translation Core of the Indiana Diabetes Research Center (2P30DK097512). The authors also thank Jacqueline Del Carmen Aquino and Lata Udari from Islet and Physiology core for their assistance with tissue analysis. We also thank Bristol Myers Squibb for providing us with the drug (BMS-986202). We thank the Stark Neuroscience Research Institute (SNRI) Biomarker Core at IU for performing GeoMx DSP transcriptomics and proteomics experiments and Dr. Emily Anderson-Baucum for her assistance in editing the text of this manuscript.

## Data Sharing Statement

Data presented in this manuscript are available from the corresponding author upon request. Genomic data generated for this manuscript has been deposited into the Genome Expression Omnibus Repository under accession number GSE262211. Reviewers can access the data using the following reviewer token: ubmhgyegtzsjnul. The proteomics data generated for this manuscript will be deposited at the time of publication.

### Declaration of Generative AI and AI-assisted Technologies in the Writing Process

During the preparation of this work the author used ChatGPT in order to organize the flow of the manuscript text. After using this tool/service, the author reviewed and edited the content as needed and takes full responsibility for the content of the publication.

### Author Contributions

FS, CEM, and DLE conceived and designed the study. FS, OB, CCL, JR, AC, SAW, ST, SD, ACdB, MIA, KO, AZ, and DS performed experiments. PK, NSC, GC, and JL performed computational analyses. FS, CEM, and DLE interpreted the data. FS and CEM wrote the manuscript. LM and PM provided islets for research. All authors provided critical revisions and edits to the manuscript. All authors read and approved of the final manuscript. CEM is the guarantor of this work. Both FS and CEM have verified the underlying data of this manuscript.

### Methods and Materials Animal Studies

Mouse studies were conducted in accordance with Animal Research: Reporting of In Vivo Experiments (ARRIVE) guidelines under protocols approved by the Indiana University (IU) School of Medicine Animal Use Committee. *RIP-LCMV-GP* mice were purchased from Jackson Laboratory and bred in-house. Eight-week-old male *RIP-LCMV-GP* mice were pre-treated with TYK2i (BMS-986202, 30 mg/kg) or vehicle (100 µL of 0.5% Methyl Cellulose-4M per mouse) by oral gavage two days prior to inoculation with LCMV. Mice continued to receive TYK2i treatment daily until the study endpoint. To monitor for T1D development, blood glucose levels were measured before inoculation and on days 1, 4, 7, 11, and 14 post-inoculation, and two consecutive blood glucose readings over 250 mg/dL established diabetes development.

Five-week-old female NOD/ShiLT mice were purchased from Jackson Laboratory and acclimated at the IU Laboratory Animal Research Center (LARC) for at least one week prior to treatment. Six-week-old female NOD mice were treated with BMS-986202 (30 mg/kg) or vehicle for 6 weeks by oral gavage and followed for up to 24 weeks to determine diabetes development. Blood glucose levels were measured weekly between ages 6-14 weeks and biweekly from 15-24 weeks of age. Two consecutive blood glucose readings over 250 mg/dL established diabetes development. To investigate insulitis, a subset of mice from each group was euthanized after 6 weeks of TYK2i treatment (12-weeks of age), and the pancreas was harvested and fixed for downstream analysis as described previously (80).

### Sex as a biological variable

In our study, we considered sex to be a biological variable. Due to the cumulative diabetes incidence in NOD female mice being 80%, compared to less than 20% in male mice (81), we exclusively used female NOD mice for our investigation. Additionally, we noted a 100% penetrance of diabetes only in male RIP-LCMV-GP mice compared to female mice. Therefore, we focused solely on male RIP-LCMV-GP mice for this study.

### Culture and treatment of human EndoC-βH1 cells and human islets

Human insulin-secreting EndoC-βH1 cells (82) (kindly provided by Dr. R. Scharfmann, Institut Cochin, University of Paris, France) were cultured in Matrigel-fibronectin-coated plates as previously described (20). The presence of mycoplasma infection was regularly controlled using the MycoAlert Mycoplasma Detection kit (Lonza).

Human islets from 8 non-diabetic organ donors were isolated in Pisa, Italy by collagenase digestion and density gradient purification with the approval of the local ethical committee and sent to Universite Libre de Bruxelles (ULB) for dispersion and experiments (20).

All experiments with EndoC-βH1 cells or human islets are shown as independent biological data (*i.e.,* considering EndoC-βH1 cells from different passages or human islets from different donors as n=1). Cells were treated with the TYK2i BMS-986165 (MedChemExpress) or BMS-986202 (a kind gift from Bristol Myers Squibb) at different concentrations ranging from 0.0003 to 3 µM and with different proinflammatory cytokines alone or in combination: IFNα (2000 U/mL), IL-1β (50 U/mL), and TNFα (1000 U/mL).

### Differentiation of iPSCs into β cells

The iPSC lines HEL46.11 and HEL115.6 were derived from fibroblasts from neonatal foreskin (83) or umbilical cord (40) obtained from healthy human donors after informed consent and approval by the Ethics Committees of the Helsinki and Uusimaa Hospital District (Helsinki, Finland) and the Erasmus Hospital (ULB, Brussels, Belgium). Cells were differentiated into pancreatic β-like cells using a 7-step protocol (40). After differentiation, stage 7 aggregates were dispersed (20) for further treatment with TYK2i and/or cytokines.

### Assessment of cell viability

Cells were stained with the DNA-binding dyes propidium iodide (PI) and Hoechst 33342 (10 µg/mL, Sigma-Aldrich) to count apoptotic cells by fluorescent microscopy, as previously described (84). This method is quantitative and has been validated by systematic comparison against electron microscopy (85) and other well-characterized methods to determine apoptosis, such as fluorometric caspase 3 & 7 assays and determination of histone-complexed DNA fragments by ELISA (86).

### mRNA extraction and quantitative real-time PCR

Cells were washed with PBS, and polyadenylated mRNA was extracted using the Dynabeads mRNA DIRECT kit (Invitrogen) and reverse transcribed with the Reverse Transcriptase Core kit (Eurogentec). Gene expression was analyzed by quantitative real-time PCR (qRT-PCR) using the SsoAdvanced Universal SYBR Green Supermix (BIO-RAD). Expression of the genes of interest was detected with the following primers (5’-3’): *HLA-ABC* fwd: CAGGAGACACGGAATGTGAA; *HLA-ABC* rev: TTATCTGGATGGTGTGAGAACC; *DDIT3(CHOP)* fwd and rev: Qiagen QuantiTect primer; *ATF3* fwd: GCTGTCACCACGTGCAGTAT; *ATF3* rev: TTTGTGTTAACGCTGGGAGA; *CXCL10* fwd: GTGGCATTCAAGGAGTACCTC; *CXCL10* rev: GCCTTCGATTCTGGATTCAG; *MX1* fwd: AGACAGGACCATCGGAATCT; *MX1* rev: GTAACCCTTCTTCAGGTGGAAC. *INS* fwd: CCAGCCGCAGCCTTTGTGA; *INS* rev: CCAGCTCCACCTGCCCCA; *PDX1* fwd: AAAGCTCACGCGTGGAAA; *PDX1* rev: GCCGTGAGATGTACTTGTTGA; *PD-L1* fwd: CCAGTCACCTCTGAACATGAA; *PD-L1* rev: ACTTGATGGTCACTGCTTGT. The quantification of amplicons was done with a standard curve, and gene expression was normalized by the expression of the reference gene *ACTB* (fwd: CTGTACGCCAACACAGTGCT; rev: GCTCAGGAGGAGCAATGATC).

### Protein extraction and Western blot analysis

Cells were lysed with 1x Laemmli Sample buffer (60 mM tris(hydroxymethyl)aminomethane pH 6.8, 10% Glycerol, 2% Sodium dodecyl sulfate, 1.5% 2-mercaptoethanol, 1.5% Dithiothreitol, and 0.005% bromophenol blue) containing phosphatase and protease inhibitors (PhosSTOP and cOmplete ULTRA tablets, mini, EASYpack protease inhibitor cocktail, Roche), and proteins were analyzed by Western blot. To evaluate STAT1 and STAT2 activation, we measured their phosphorylation using a rabbit anti-phospho-STAT1 (cs9167, Cell Signaling, 1:1000) and a rabbit anti-phospho-STAT2 antibody (cs8841, Cell Signaling, 1:4200). Immunoreactive bands were detected using ChemiDoc XRS+ (Bio-Rad) after incubation with a secondary anti-rabbit antibody coupled with the horseradish peroxidase (HRP, 1:5000) and exposure to the SuperSignal West Femto chemiluminescent substrate (Thermo Fisher Scientific). Densitometric quantification of the bands was performed with ImageLab software (Bio-Rad). Data were normalized for the expression of the housekeeping protein β-actin (cs4967, Cell Signaling, 1:5000).

### Generation of EndoC-βH1 cells expressing HLA-A2 and T cell activation assays

EndoC-βH1 cells expressing HLA-A2 were generated by lentivirus transduction (87) using a lentiviral vector containing HLA-A02:01 under the elongation factor 1α (EF1α) promotor. These cells were obtained from R.J. Lebbink (Medical Microbiology, University Medical Center Utrecht, Utrecht, the Netherlands) (88). Target cells (i.e. EndoC-βH1 cells expressing HLA-A2) were harvested and cocultured with cytotoxic lymphocytes (CTLs) specific for an epitope located within the pre-proinsulin signal peptide (ALWGPDPAAA) (89) at an effector/target ratio of 2:1 in the presence of mouse anti-human CD107a-FITC (11-1079-42; Thermo Fisher Scientific). Cocultures between CTLs and EndoC-βH1 cells expressing HLA-A2 were performed at 37 °C for 1.5 h in Iscove’s Modified Dlbecco’s Medium (IMDM) supplemented with 10% human serum and 25 U/mL IL-2 (Novartis), as described previously (90,91). T cell activation is shown as relative degranulation, estimated by calculating the percent change of the absolute degranulation of the treated samples compared to their respective controls.

### Flow cytometry

Spleen, PLNs, and blood were collected from TYK2i-or vehicle-treated *RIP-LCMV-GP* mice. Spleen and PLNs were homogenized and passed through a 70 μm cell strainer (BD Falcon, San Jose, CA) to obtain a single-cell suspension. Cells were incubated with mouse anti-CD16/32 (BioLegend) to block Fc receptors and then surface-stained with fluorochrome-conjugated monoclonal antibodies for 20 mins at 4 °C followed by PBS wash. Live/dead staining was performed using Zombie aqua (1:1000 dilution, BioLegend) for 20 mins at room temperature to exclude dead cells. Multivariable flow cytometry was performed using anti-mouse monoclonal antibodies: CD3-FITC (17A2; ThermoFisher), CD19-A700 (1D3), CD11b-BV650 (M1/70), CD11c-APC (N418), MHCII-PerCPCy5.5 (M5/114.15.2), F4/80-PE (BM8), Ly6c-PECy7 (HK1.4), CD49b-APCCy7 (DX5), CD4-A700 (GK1.5), CD8a-PerCPCy5.5. (53-6.7), CD25-BV650 (PC61), CD44-APCCy7 (IM7), CD62L-PE (MEL-14), CD69-PECy7 (H1.2F3), PD1-BV605 (29F.1A12), and CXCR3-BV421 (S18001A). All the antibodies except anti-CD3 FITC were from BioLegend.

Single cell suspensions were permeabilized for intracellular staining using FoxP3 fix/perm buffer (ThermoFisher) at 4 °C for 20 mins, followed by a wash with permeabilization buffer. Samples were then stained with mouse anti-FoxP3-APC (FJK-16s; ThermoFisher) for 30 mins at 4 °C, followed by PBS wash. Cell suspensions were evaluated on an Attune NxT flow cytometer (ThermoFisher), and all analysis was performed in FCS Express (Version 7, De Novo Software).

### Immunohistochemistry

Pancreatic tissues were harvested and fixed with 4% PFA overnight at 4 °C and tissues were processed and embedded in paraffin blocks as described previously (80). Tissue sections were stained with rabbit anti-insulin antibody (Cell Signaling) and counterstained with DAB peroxidase anti-rabbit IgG (Vector Lab). Images were acquired using a Zeiss slide scanner (Zeiss, Germany), and insulitis scoring was performed on 5 slides, each 30 µm apart, from 9 mice/group as previously described (80). Immunofluorescence studies were performed according to the protocol described previously (46) using the following antibodies: guinea pig anti-insulin (Dako), mouse anti-PD-L1 (Proteintech), and rabbit anti-CXCL10 (Invitrogen). Signals were detected by counter staining with the following antibodies: goat anti-guinea pig (1:400; Invitrogen), goat anti-rabbit (1:200; Invitrogen), and goat anti-mouse (1:200; Invitrogen). All sections were counterstained with DAPI to identify nuclei. Images were obtained with a Zeiss LSM 800 confocal microscopy (Carl Zeiss, Germany), the fluorescence intensities were quantified using Image J, and the corrected fluorescence intensities were presented, as described previously (46).

### Single-molecule fluorescence *in situ* hybridization

PFA fixed and paraffin-embedded tissue sections cut under RNase-free conditions were pre-treated and processed as described in our previous studies (92,93). Fluorescence *in situ* hybridization was performed using an RNAscope Multiplex Fluorescent Detection kit (RNAscope) according to the manufacturer’s protocol. Briefly, a combination of two probes targeting mouse genes (*CD274* and *Mx1* or *Cxcl10* and *Stat1* at day 3 and day 14; and *CD274* and *Cxcl10* or *Mx1* and *Stat1 at day 7)* was hybridized for 2 h, and the signals were detected using secondary TSA fluoroprobes using the cyclic detection method for two different channels with co-staining for insulin and DAPI. The hybridized sections were mounted using ProLong Gold Antifade mounting media (Thermo Fisher Scientific) and imaged using a Zeiss LSM800 microscope (Carl Zeiss, Germany) attached to an Airyscan detector.

### smFISH analysis

The processing, analysis, and quantification of smFISH images was performed similar to our previous work (94), which involves a series of image detection algorithms and data processing tools. Initially, the multi-channel images were segregated into four distinct channels, including the nucleus channel, the insulin protein channel, and two mRNA channels. We used a combination of manual segmentation and the implementation of the Segment Anything Model (SAM) (95) to identify the nucleus from the DAPI channel. Post-processing of the SAM-segmented nucleus masks involved the removal of duplicate masks and the elimination of segmented objects with areas smaller than 3000 pixels (approximately 2.7 μm^2^). Subsequently, the nuclear boundary was extended radially to predict the cell boundary. The identification of β cells was achieved by overlaying the segmented cells with the insulin protein channel, allowing for the recording of insulin protein intensity for each cell. To further analyze the β and the non-β cell populations, a histogram was plotted based on the resulting insulin protein intensity per cell. Savitzky-Golay filter was applied to smooth the histogram by yielding two distinct cell populations. By identifying the local minima in the smoothed histogram, the separation between distributions was determined, establishing the insulin intensity threshold for β cell identification.

To detect the individual mRNA particles from the two mRNA channels, the Laplacian of Gaussian (LoG) algorithm was utilized (96). This algorithm provides estimated locations of mRNA particles, facilitating subcellular expression analysis. In some instances, multiple mRNA particles may exhibit intense brightness due to overlapping at close proximity, showing a single bright focus on smFISH images. To address this issue, the number of mRNAs at a bright focus was deduced by normalizing the focus intensity of the single mRNA intensity (97). The intensity of a single mRNA was set by binning mRNA particles based on intensity and identifying the intensity bin containing the majority of mRNAs. Finally, outliers caused by autofluorescence were further filtered out using Tukey’s fence (98) to prevent overcounting of RNAs.

### Nanostring GeoMx DSP spatial transcriptomics

Mouse pancreas sections (5 µm) were fixed, sectioned, and mounted on positively charged slides. The slides were deparaffinized, heated in antigen retrieval solution (Thermo Fisher Scientific) at 100 °C for 20 min, and treated with 1 µg/mL proteinase K at 37 °C for 15 min. Mouse transcriptomics was visualized using the GeoMx mouse whole transcriptome atlas (NanoString Technologies, GMX-RNA-NGS-MsWTA). An overnight *in situ* hybridization was performed with a mouse WTA probe, and the next day, the slides were washed twice at 37 °C for 25 min with 50% formamide/2X SSC buffer to remove unbound probes. The slides were then washed twice with 2X SSC buffer for 2 minutes at RT and blocked with blocking buffer containing Buffer W (Nanostring) with 1% Fc-Receptor blocker (Milteny) and 5% Normal Donkey Serum (Jackson Laboratories) as described previously (99). The slides were incubated with guinea pig anti-insulin (Dako), 1:200 rabbit anti-CD3 (Abcam), mouse anti-CD68 (Santa Cruz). The slides were washed twice with 2xSSC and incubated with secondary antibodies (1:400 goat-anti-guinea pig Alexa-488; 1:200 goat-anti rabbit Alexa-594 and 1:200 goat-anti mouse Alexa-647) for 30 minutes. Then, the slides were washed four times with 2xSSC for 3 minutes and counter-stained with 1:20,000 Syto83 (Invitrogen) for 1 h at room temperature. Stained slides were loaded onto the GeoMx instrument and scanned. After visual inspection using the GeoMx Digital Spatial Profiler (DSP; NanoString Technologies), geometrical ROIs (100 μm diameter) with different histologies were selected for oligonucleotide collection. Photocleaved oligonucleotides from each spatially resolved ROI were sequenced with the Illumina NextSeq500 sequencer (Illumina). The GeoMx libraries were prepared according to the GeoMx DSP NGS Readout User Manual. Briefly, for each plate, 4 μl of a GeoMx DSP sample was used for amplification. Each GeoMx DSP sample in a well was uniquely indexed with dual indexes. 4 μL of each resulting PCR product, including the NTC negative control, were pooled and purified with AMPure XP beads (Beckman Coulter). The final libraries were qualified and quantified using TapeStation and Qubit, respectively. The libraries were sequenced on an Illumina NextSeq 500 to generate 27 bp paired-end sequence reads.

### Spatial transcriptomics data analysis

Spatial transcriptomics analysis was conducted in R studio v4.2.1 using the GeomxTools 3.4.0 package (100). The GeoMx whole mouse transcriptome atlas was used as the probe assay metadata. Sequencing quality was assessed using the default quality control (QC) parameter cut-offs, namely: minimum number of reads (1000); minimum percentage of reads trimmed (80%), stitched (80%), and aligned (80%); sequencing saturation (50%); negative control counts (10); maximum counts observed in no treatment control (NTC) well (1000); minimum number of nuclei estimated (100); and minimum segment area (5000). Any ROI that did not satisfy the above criteria was removed as part of QC filtering. Following segment QC, probe QC was also performed to remove any outlier either globally or locally before generating gene-level count data. The limit of quantification was determined per segment using the value recommended in the workflow, and segments with less than 1-5% of genes detected were removed. Furthermore, only genes that were detected in at least 1% of segments were retained. The data was normalized using the quartile 3 method and log transformed. Samples were clustered based on overall gene expression using either UMAP (umap package) or tSNE (Rtsne package). Genes with a high coefficient of variation were plotted using unsupervised hierarchical clustering displayed as a heatmap. For each organ, a linear mixed model without random slope was used to identify differentially expressed genes between vehicle and treatment groups for both the 3- and 7-day timepoints. Genes with linear scale fold change ≥ 1.5 and p-value < 0.05 were considered as differentially expressed. These differentially expressed genes were visualized using volcano plots, and the top ten differentially expressed genes from each region were visualized using heatmaps (pheatmap function from gplots package). All graphics were generated using ggplot2 unless otherwise stated. Pathway analysis for each time point from each organ was performed using the Qiagen Ingenuity pathway analysis tool (QIAGEN Inc., https://digitalinsights.qiagen.com/IPA). Pathways with a z-score of at least 1 and p-value < 0.05 were considered for further interpretation. Spatial deconvolution for each organ and every time point was performed using the SpatialDecon package (http://www.bioconductor.org/packages/release/bioc/manuals/SpatialDecon/man/SpatialDecon.pdf). Normalized values obtained from the above workflow were used as the input for the expression matrix. Adult mouse pancreas (Pancreas_MCA) and adult mouse spleen (Spleen_MCA) were used as cell matrices for determining the cell populations in the pancreas and spleen, respectively. In addition, the mouse immune profile (ImmuneAtlas_ImmGene) was used as the cell matrix for estimating immune cell abundance in the pancreas, spleen, and PLN. The cell matrices were downloaded from (101,102). Basic deconvolution was performed following the workflow provided in the R package. For immune cell populations, the cells were merged based on the lineage. Beta values were used as the input for heatmap (pheatmap function from gplots package) and abundance plots. Any population with a zero value was removed from the graphs. Abundance plots were illustrated as stacked bar charts using GraphPad Prism, and the difference between vehicle and treatment groups was compared using a student’s t-test (two-tailed). Cell populations with p-value < 0.05 were considered significant.

#### Spatial proteomics using the NanoString GeoMx spatial profiling platform

Pancreatic sections were deparaffinized and rehydrated, similar to the transcriptomic steps described above. Next, antigen retrieval was performed with 1X citrate buffer pH 6.0 in a preheated pressure cooker for 15 min. The slides were then stained with anti-insulin (AlexaFluor 488, 1:50; R&D Systems), anti-B220 (AlexaFluor 594, 1:150;), and anti-CD3 (AlexaFluor 647, 1:200; Bio-Rad) antibodies. In addition, DSP antibody barcode collections including immune cell core profiling, immune cell activation, and immune cell typing (nanoString) were added to the slides and incubated overnight at 4 °C. Nuclei were identified by staining with Syto83 fluorescent dye (0.2 µm concentration, Fluorescence 532, Thermo Fisher Scientific). The corresponding ROIs from the protein slide were selected and collected in a collection plate using a similar method as the RNA profiling. The aspirates were then reacted directly with Probe R and Probe U (Nanostring Technologies), which carry the fluorescent reporters and can be used for fluorescence counting with the nCounter Max Analysis system (Nanostring Technologies, Inc).

### GeoMx DSP proteomic data analysis

Protein expression counts from the nCounter run were retrieved and uploaded into the GeoMx DSP analysis suite (Nanostring, version 3.0.109), and the nCounter read counts were normalized using a geometric mean of the signal-to-noise ratio (SNR) of IgG negative controls as described previously (99,103). The normalized read counts are presented as Log_2_ of SNR, and the difference was presented with false discovery rate (FDR).

### Statistical analysis and reproducibility

The quantitative polymerase chain reaction (qPCR) and Western blot analyses were conducted on datasets comprising 3-6 biological replicates. To determine statistical significance, we employed a one-way analysis of variance (ANOVA) with subsequent Bonferroni correction for multiple comparisons. For the analysis of diabetes onset data, we applied the Log-rank (Mantel-Cox) test, while insulitis scoring was analyzed using a one-sample t-test. The Mann-Whitney test was used to evaluate variations in mRNA and protein expression between vehicle- and TYK2i-treated *RIP-LCMV-GP* and NOD mice. All data analyses were performed utilizing GraphPad Prism, version 10. The statistical analysis for spatial transcriptomics and proteomics data are discussed above in their respective methods section.

---

**Supplemental Figure 1.**
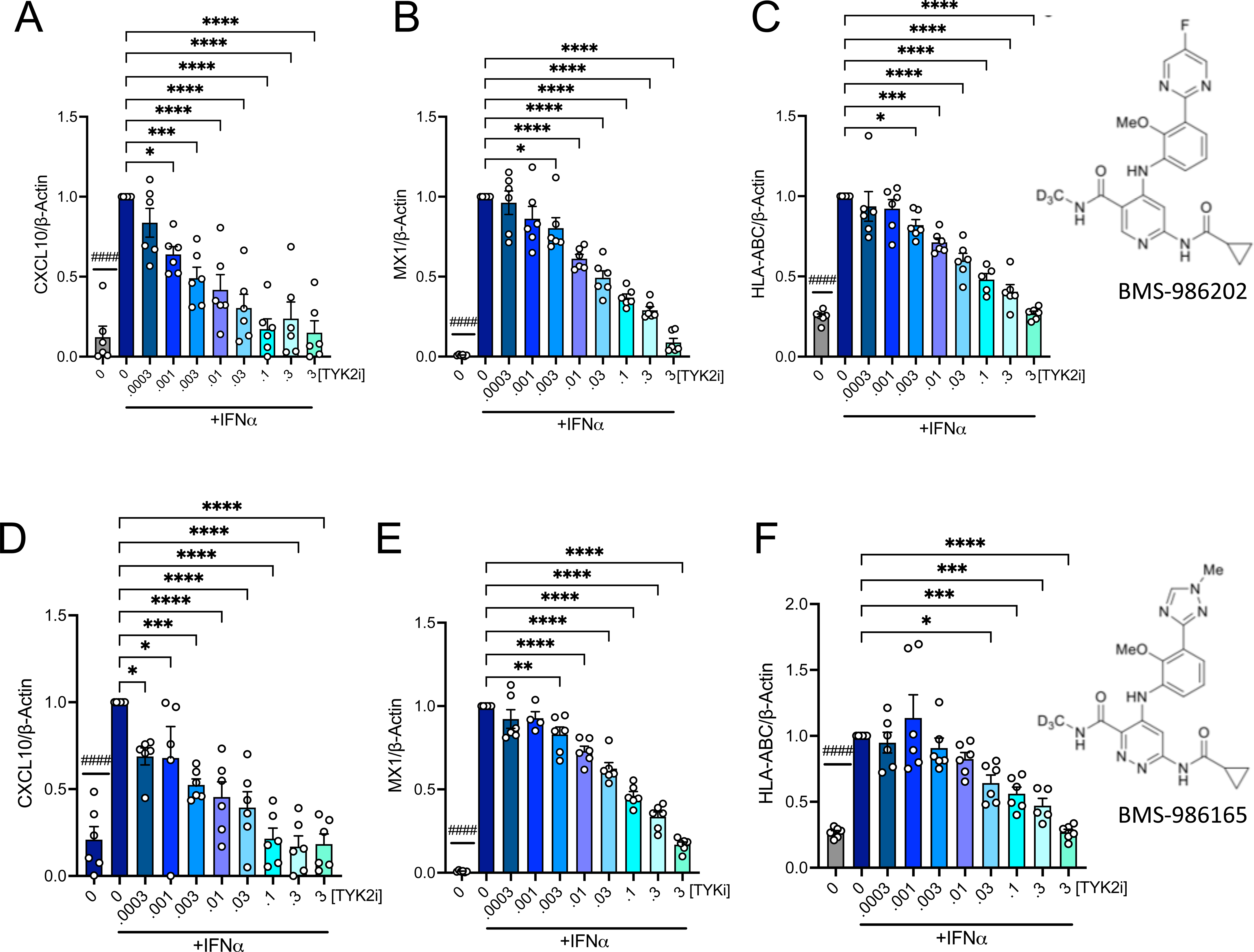

**Supplemental Figure 1.**
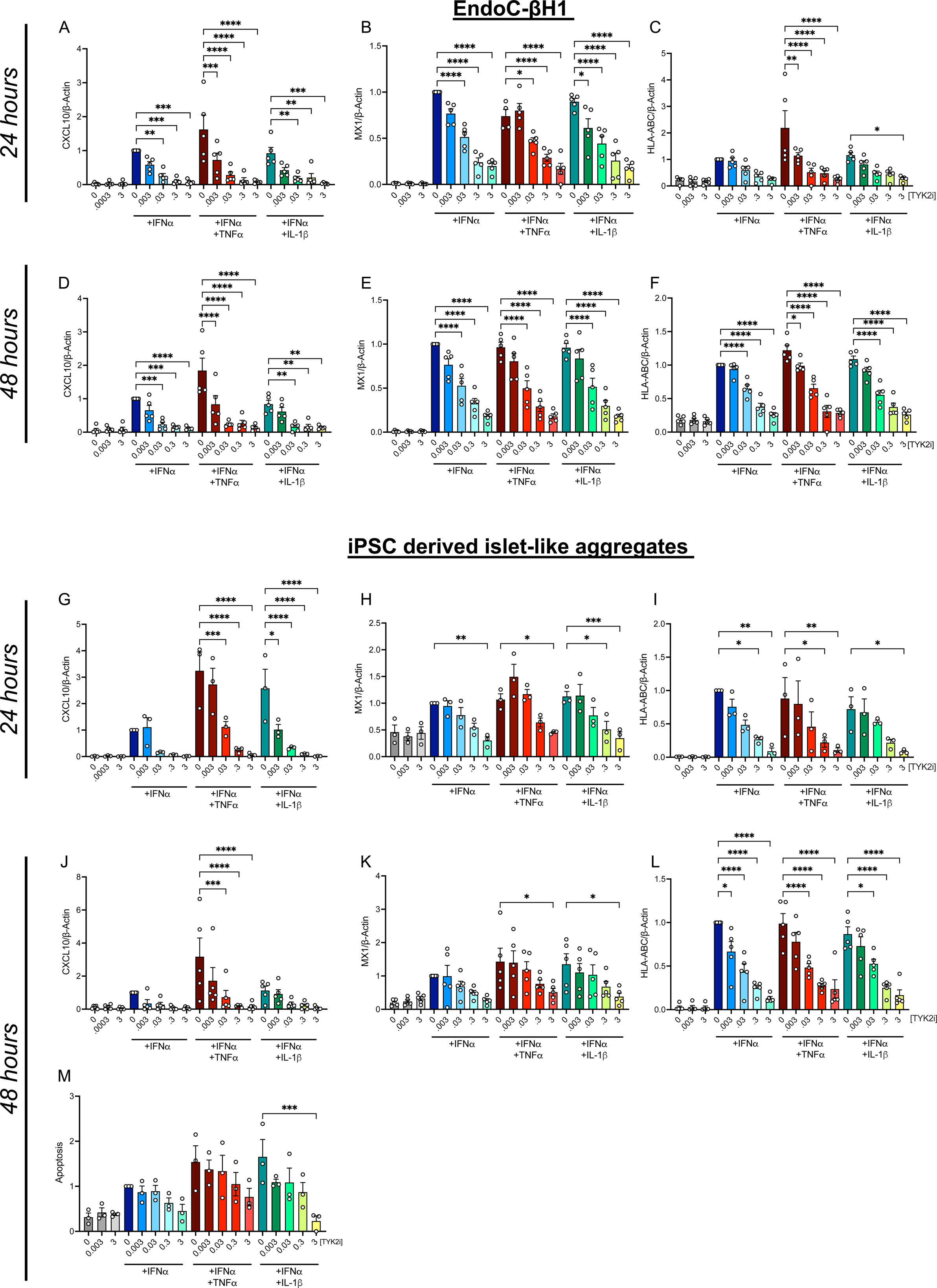

**Supplemental Figure 1.**
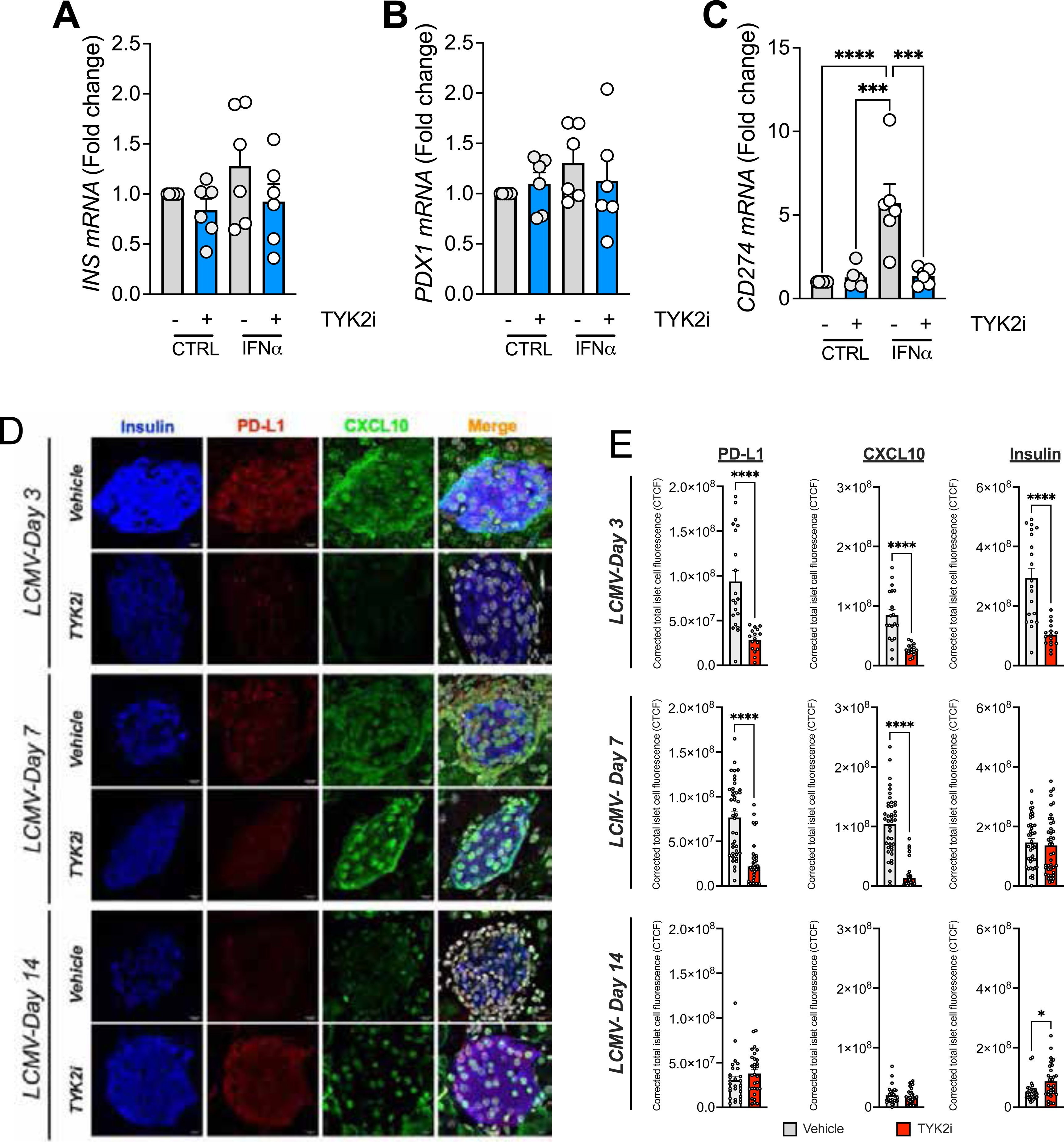

**Supplemental Figure 1.**
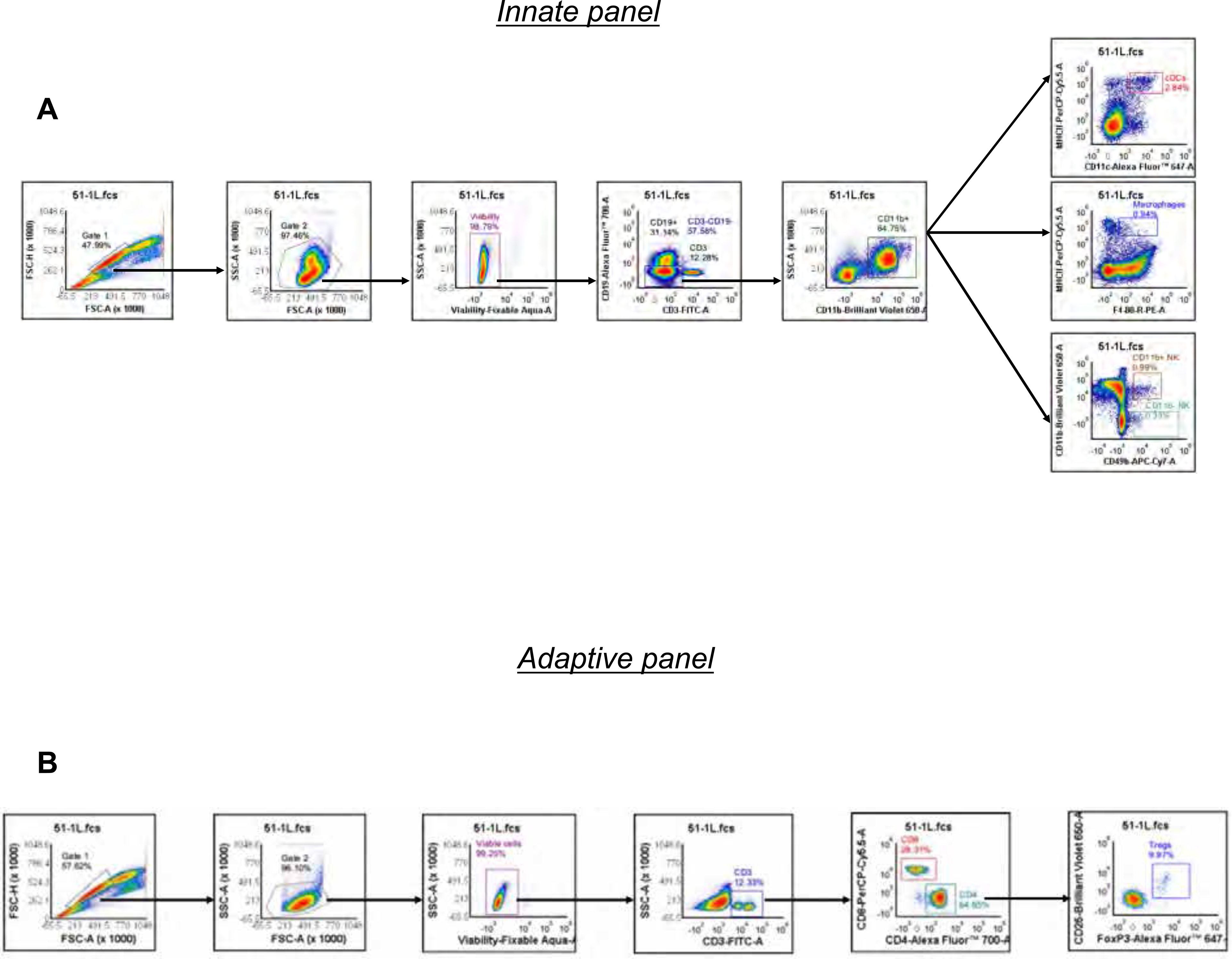

**Supplemental Figure 1.**
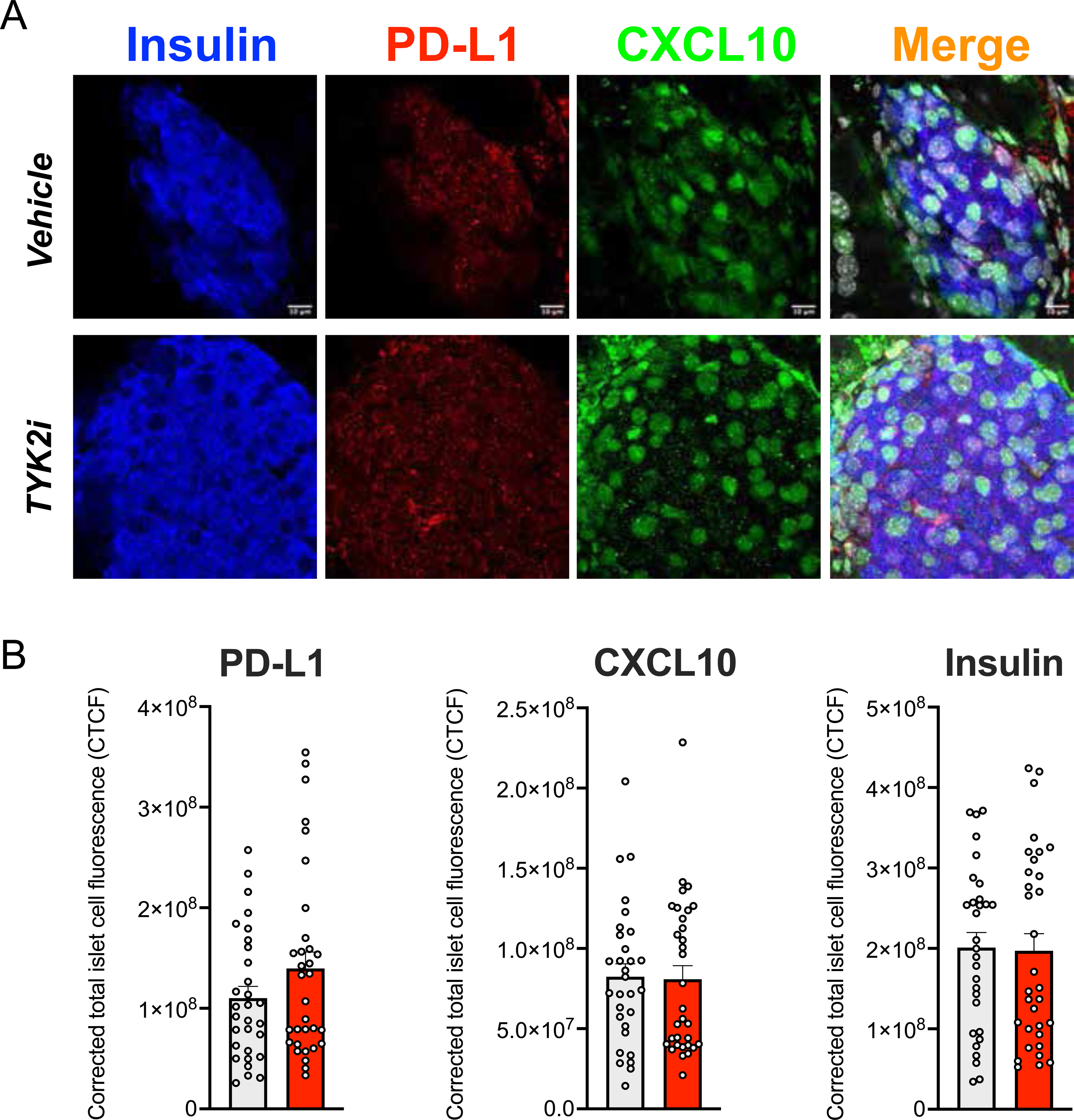

**Supplemental Figure 1.**
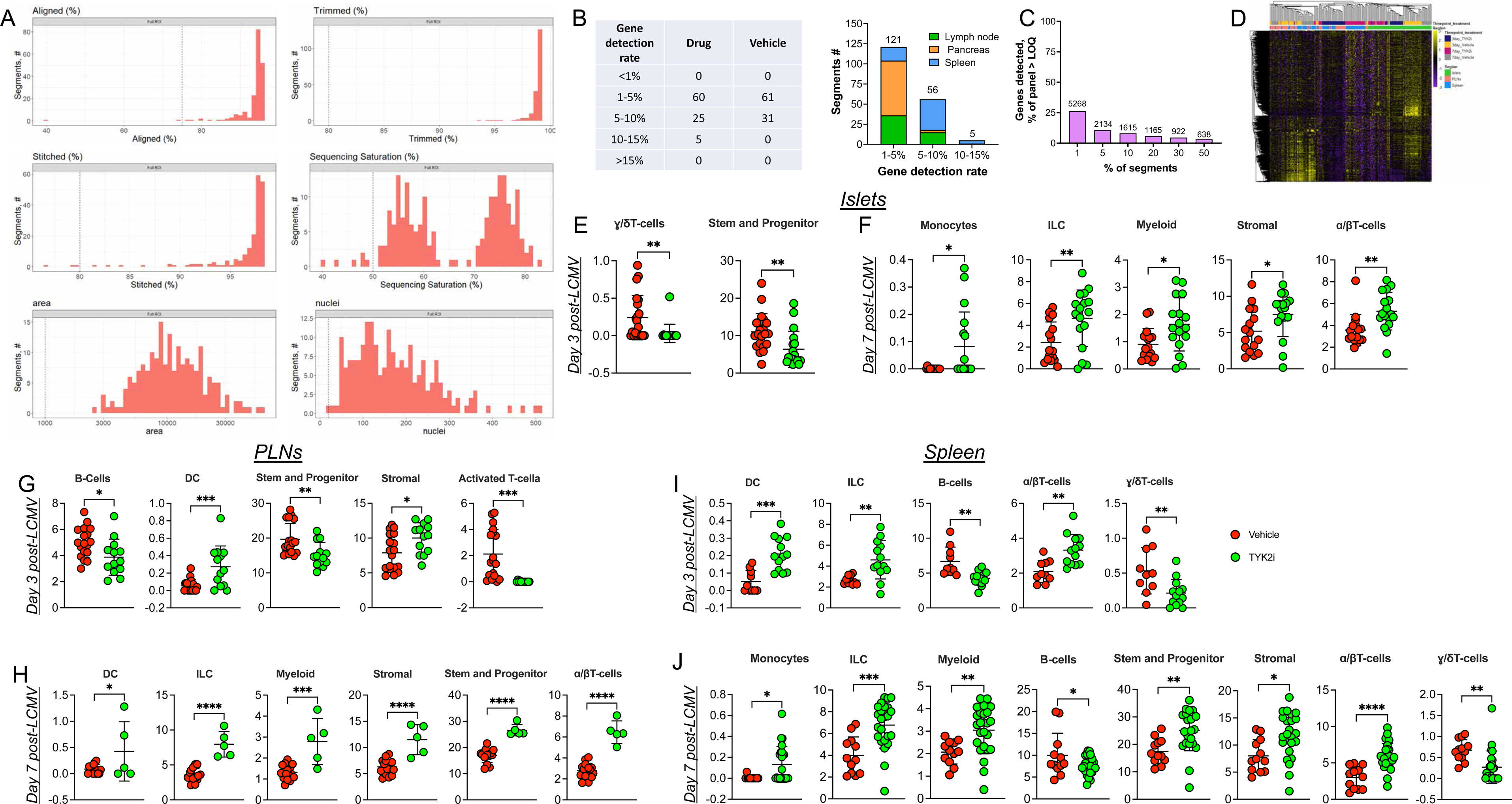

**Supplemental Figure 1.**
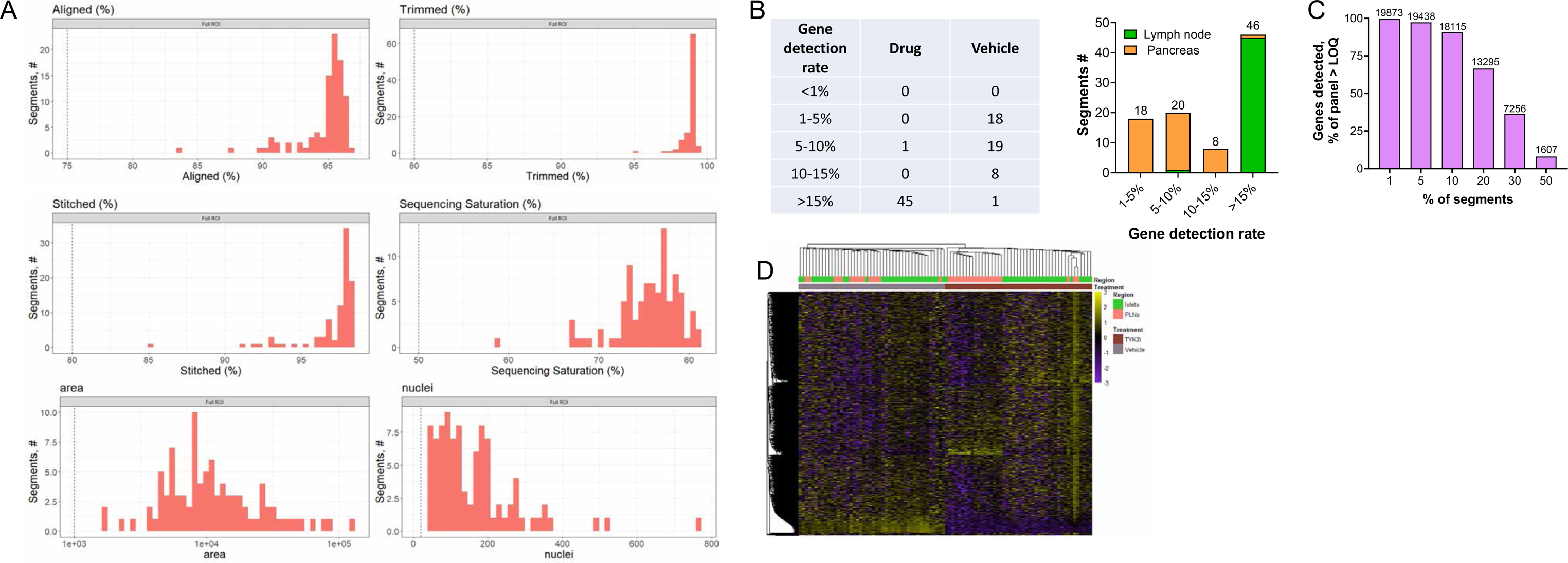

**Supplemental Figure 1.**
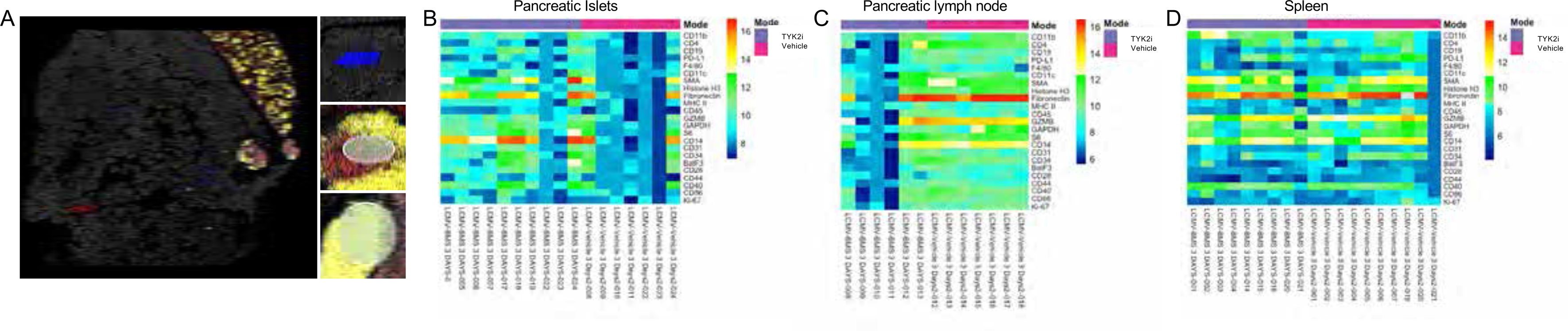

## Notes

### Competing Interest Statement

CEM has served on advisory boards related to T1D research clinical trial initiatives: Provention Bio, Dompe Pharmaceuticals, Isla Technologies, MaiCell Technologies, Avotres, and DiogenX. CEM has patent (16/291,668) Extracellular Vesicle Ribonucleic Acid (RNA) Cargo as a Biomarker of Hyperglycaemia and Type 1 Diabetes and CEM and FS have a provisional patent (63/285,765) Biomarker for Type 1 Diabetes (PDIA1 as a biomarker of β cell stress). These activities have not dealt directly with topics covered in this manuscript.

### Summary of Updates

Revision to reflect the most current version of the manuscript.

