## Supplementary material for "Pharmacological inhibition of tyrosine protein-kinase 2 reduces islet inflammation and delays type 1 diabetes onset in mice": Human Islet Checklist

**Checklist for reporting human islet preparations used in research**

Adapted from Hart NJ, Powers AC (2018) Progress, challenges, and suggestions for using human islets to understand islet biology and human diabetes. Diabetologia <https://doi.org/10.1007/s00125-018-4772-2>

| **Islet preparation** | **1** | **2** | **3** | **4** | **5** | **6** | **7** | **8** |
| --- | --- | --- | --- | --- | --- | --- | --- | --- |
| **MANDATORY INFORMATION** | | | | | | | | |
| Unique identifier | 26/115 | 26/117 | 26/128 | 26/133 | 26/134 | 27/7 | 27/14 | 27/17 |
| Donor age (years) | 86 | 81 | 62 | 63 | 65 | 76 | 86 | 73 |
| Donor sex (M/F) | F | M | F | M | M | F | M | F |
| Donor BMI (kg/m^2^) | 27.1 | 27.1 | 23.5 | 29.4 | 27.8 | 23.9 | 26.1 | 25.0 |
| Intensive Care Unit glycemia (mg/dl) | 196 | 187 | 163 | 73 | 252 | 177 | 149 | 177 |
| Origin/source of islets^b^ | Cisanello University Hospital, Pisa | Cisanello University Hospital, Pisa | Cisanello University Hospital, Pisa | Cisanello University Hospital, Pisa | Cisanello University Hospital, Pisa | Cisanello University Hospital, Pisa | Cisanello University Hospital, Pisa | Cisanello University Hospital, Pisa |
| Islet isolation centre | Cisanello University Hospital, Pisa | Cisanello University Hospital, Pisa | Cisanello University Hospital, Pisa | Cisanello University Hospital, Pisa | Cisanello University Hospital, Pisa | Cisanello University Hospital, Pisa | Cisanello University Hospital, Pisa | Cisanello University Hospital, Pisa |
| Donor history of diabetes? (Yes/No) | No | No | No | No | No | No | No | No |
| **If Yes, complete the next two lines if this information is available** | | | | | | | | |
| Diabetes duration (years) |  |  |  |  |  |  |  |  |
| Glucose-lowering therapy at time of death^c^ |  |  |  |  |  |  |  |  |
| **RECOMMENDED INFORMATION** | | | | | | | | |
| Donor cause of death | Cardiovascular disease | Cardiovascular disease | Cardiovascular disease | Cardiovascular disease | Cardiovascular disease | Cardiovascular disease | Cardiovascular disease | Cardiovascular disease |
| Warm ischaemia time (h) |  |  |  |  |  |  |  |  |
| Cold ischaemia time (h) | 15 | 15 | 15 | 18 | 14 | 18 | 13 | 17 |
| Estimated purity (%) |  |  |  |  |  |  |  |  |
| Estimated viability (%) |  |  |  |  |  |  |  |  |
| Total culture time (h)^d^ |  |  |  |  |  |  |  |  |
| Glucose-stimulated insulin secretion or other functional measurement^e^ | Glucose-stimulated insulin secretion | Glucose-stimulated insulin secretion | Glucose-stimulated insulin secretion | Glucose-stimulated insulin secretion | Glucose-stimulated insulin secretion | Glucose-stimulated insulin secretion | Glucose-stimulated insulin secretion | Glucose-stimulated insulin secretion |
| Handpicked to purity? (Yes/No) | Yes | Yes | Yes | Yes | Yes | Yes | Yes | Yes |
| Additional notes |  |  |  |  |  |  |  |  |

^a^If you have used more than eight islet preparations, please complete additional forms as necessary

^b^For example, IIDP, ECIT, Alberta IsletCore

^c^Please specify the therapy/therapies

^d^Time of islet culture at the isolation centre, during shipment and at the receiving laboratory

^e^Please specify the test and the results
