## Supplemental Figure Legends for "Pharmacological inhibition of tyrosine protein-kinase 2 reduces islet inflammation and delays type 1 diabetes onset in mice"

**Supplemental Figure 1. TYK2 inhibitors BMS-986202 and BMS-986165 alleviate IFNα-induced gene expression in EndoC-βH1 cells.**

(**A-F**) Cells were pre-treated for 2 h with the indicated concentrations (expressed in mM) of (**A-C**) BMS-986165 or (**D-F**) BMS-986202 and exposed to IFNα (2000 U/mL) in the continued presence of the inhibitors for 24 h. mRNA expression of (**A,D**) *CXCL10*, (**B,E**) *MX1*, and (**C,F**) *HLA-ABC* was analyzed by qPCR. Data was normalized to β-actin expression level and expressed as fold changed compared to cells exposed to IFNα alone. Data are presented as mean ± SEM for 6 independent experiments with individual data indicated; ####*p*<0.0001 vs no inhibitor and IFNα and **p*<0.05, ***p*<0.005, ****p*<0.001, and *****p*<0.0001 vs no inhibitor and IFNα; one-way ANOVA followed by Bonferroni correction for multiple comparisons.

**Supplemental Figure 2. TYK2 inhibition suppresses cytokine-induced gene expression in human β cells and iPSC-derived β-like cells.**

(**A-F**) EndoC-βH1 cells were pre-treated for 2 h with the indicated concentrations (expressed in mM) of BMS-986202 prior to exposure to IFNα (2000 U/mL), IFNα (2000 U/mL) + TNFα (1000 U/ml), or IFNα (2000 U/mL) + IL-1β (50 U/mL) in the continued presence of the TYK2i for **(A-C)** 24 h or **(D-F)** 48 h. mRNA expression of (**A,D**) *CXCL10*, (**B,E**) *MX1*, and (**C,F**) *HLA-ABC* was analyzed by qPCR. (**G-I**) Control iPSCs (HEL115.6) were differentiated into islet-like cells using a previously described protocol (41) and pre-treated for 2 h with the indicated concentrations (expressed in mM) of BMS-986202 prior to exposure to IFNα (2000 U/mL), IFNα (2000 U/mL) + TNFα (1000 U/ml), or IFNα (2000 U/mL) + IL-1β (50 U/mL) in the continued presence of the TYK2i for (G-I) 24 h or (J-M) 48 h. mRNA expression of (**G, J**) *CXCL10*, (**H, K**) *MX1*, and (**I, L**) *HLA-ABC* was analyzed by qPCR. Islet-like aggregates were dissociated and seeded in 8-well chamber slides. After 48 h, cells were pre-treated for 2 h with the indicated concentrations (expressed in mM) in BMS986202 and exposed to IFNα (2000 U/mL), IFNα (2000 U/mL) + TNFα (1000 U/mL), or IFNα (2000 U/mL) + IL-1β (50 U/mL) in the continued presence of inhibitor for 48 h. (**M**) Apoptotic cells were identified by Hoechst 3342 and propidium iodide staining. For qPCR analysis, data was normalized for β-actin expression and expressed as fold change from cells exposed to IFNα alone. Data are presented as means ± SEM. n= 5 for EndoC-βH1 cells. For iPSC cells n=3 for 24 h and n=6 for 48 h timepoints. Independent experiments with individual data are indicated. For apoptosis studies in iPSC derived islet-like aggregates, data are expressed as fold change compared to cells exposed to IFNα and are presented as mean ± SEM for 3 independent experiments with individual data indicated. ####*p*<0.0001 vs no inhibitor and IFNα (2000 U/mL) and **p*<0.05, ***p*<0.005, ****p*<0.001, *****p*<0.0001 vs no inhibitor and with indicated cytokines, one-way ANOVA followed by Bonferroni correction for multiple comparisons.

**Supplemental Figure 3. TYK2 inhibition does not affect mRNA expression of β cell-specific genes but inhibits *CXCL10* and *CD274* expression.**

(**A-C**) Human islets were pretreated with DMSO or BMS-986165 (0.3 µM) for 2 h and then co-incubated with or without IFNα (2000 IU/mL) for 24 h. qPCR analysis of (**A**) *INS*, (**B**) *PDX1*, and (**C**) *CD274*. Results are presented as mean ± SEM for 5 independent experiments with individual data indicated. ***p<0.001, ****p<0.0001, one-way ANOVA followed by Bonferroni correction for multiple comparisons. (**D-E**) Pancreatic tissue was harvested from vehicle- and TYK2i-treated *RIP-LCMV-*GP mice on days 3, 7, and 14 post-inoculations. (**D**) Representative immunofluorescence images of PD-L1 (red) and CXCL10 (green) in pancreatic islets at 3, 7, and 14 days post-inoculation. (**E**) Quantification of immunofluorescence signals presented as corrected cellular fluorescence intensities. Grey and red bars indicate vehicle- and TYK2i-treated groups, respectively. Data are expressed as mean ± SEM with individual data presented, n=5-7 mice per condition and 4-7 islets per section; **p*≤0.01, *****p*≤0.0001, Statistical significance was determined by Mann-Whitney test.

**Supplemental Figure 4. Gating strategy for flow cytometry analysis from *RIP-LCMV-GP* mice.**

Representative images of scatter plots showing immune cell markers used to determine the abundance of (**A**) innate and (**B**) adaptive immune cells in the blood, PLN, and spleen of *RIP-LCMV-GP* mice.

**Supplemental Figure 5. TYK2 inhibition does not affect IFNα-induced PD-L1 and CXCL10 protein expression in islets of prediabetic NOD mice.**

Pancreas tissue was harvested from vehicle- or TYK2i-treated NOD mice at 13 weeks of age. (**A-B**) Immunofluorescent labeling for the detection of insulin, PD-L1, and CXCL10 in pancreas tissue sections. (**A**) Representative confocal images and (**B**) corrected total cellular fluorescence intensities from islets of vehicle- and TYK2i-treated NOD mice. Grey and red bars indicate measures in tissues from vehicle- or TYK2i-treated mice, respectively. Data are expressed as mean ± SEM with individual data presented, n=4-6 mice per condition; **p*≤0.05, ***p*≤0.01, ****p*≤0.001, statistical significance was determined by Mann-Whitney test.

**Supplemental Figure 6. Quality control and normalization strategies used in spatial whole transcriptome analysis of *RIP-LCMV-GP* mice.**

(**A**) Percentage of aligned sequencing read counts at every step of sequencing and sequencing saturation. (**B**) Gene detection rate across different cutoff ratios and (**C**) number of genes detected across different filtering criteria. (**D**) Heatmap showing gene clustering using genes with a high coefficient of variation. (**E-J**) Abundance of immune cell populations was identified by deconvolution analysis of WTA using immune cell matrix pipeline at day 3 and 7 post-inoculation from (**E-F**) islets, (**G-H**) PLN,s and (**I-J**) spleen of the vehicle and TYK2i-treated *RIP-LCMV-GP* mice.

**Supplemental Figure 7. Quality control and normalization strategies used in spatial whole transcriptome analysis of NOD mouse.**

(**A**) Percentage of aligned sequencing read counts at every step of sequencing and sequencing saturation. (**B**) Gene detection rate across different cutoff ratios and (**C**) number of genes detected across different filtering criteria. (**D**) Heatmap showing gene clustering using genes with a high coefficient of variation from the islets and PLN of the vehicle- and TYK2-treated NOD mice.

**Supplemental Figure 8. Spatial proteomics of islets, PLN, and spleen of *RIP-LCMV-GP* mice treated with vehicle or TYK2 inhibitor at day 3.**

(**A**) Representative images of islets, PLN, and spleen of vehicle- or TYK2i-treated *RIP-LCMV-GP* mice at day 3 post-inoculation labeled for CD3 (red), PTPRC (yellow), insulin (blue), and sytox83 (grey). (**B-D**) Heatmap showing the overall expression of immune cell typing and immune cell activation markers from selected regions of interest (ROIs) from (**B**) islets, (**C**) PLN, and (**D**) spleen; n=3 mice/group. Data were normalized to the geometric mean of the IgG negative control, and the significantly expressed proteins were presented as Log_2_ of signal-to-noise ratio (SNR).
